# A haplotype-based evolutionary history of barley domestication

**DOI:** 10.1101/2024.12.18.628695

**Authors:** Yu Guo, Murukarthick Jayakodi, Axel Himmelbach, Erez Ben-Yosef, Uri Davidovich, Michal David, Anat Hartmann-Shenkman, Mordechai Kislev, Tzion Fahima, Verena J. Schuenemann, Ella Reiter, Johannes Krause, Brian J. Steffenson, Nils Stein, Ehud Weiss, Martin Mascher

## Abstract

Barley is an old crop with a complex history. Its evolution and the molecular basis of domestication have been intensely studied. This research has ruled out a single origin of the crop and motivated a model of mosaic genomics ancestry. As more and better genome sequences have become available, this concept can be refined: where do the building blocks of the mosaic, that is, the haplotypes come from? Were all cultivated barleys created equal or did some wild barley populations contribute more to some domesticated lineages? To answer these and other questions, we pursue a haplotype-based approach to connect diversity and population structure in wild and domesticated barley. We sequence the genomes of 628 genebank accessions and 23 archaeological specimens. Using these data, we infer the spatiotemporal origins of haplotypes and map the contributions of different wild barley populations either during the initial phase of domestication or through later gene flow. Ancient DNA sequences corroborate our genome-wide analysis in present-day samples. Our results indicate that an early domesticated founder population formed in the Fertile Crescent during an extended period of pre-domestication cultivation. A practical implication of our findings is that the high haplotype differentiation between barley populations, which arose possibly without, and most likely on top of, selective forces, complicates the mapping of adaptive loci.

---

Barley (*Hordeum vulgare*) is an old crop. It is mentioned in some of the earliest records of human writing (3100 BCE)^1^. By that time, plant cultivation was older than written language is now. Much of what we know about the early stages of the domestication and dispersal of barley and other crops comes from archaeological specimens, the earliest dated to 10,000 years before the present (BP)^2,3^. These are mainly charred grains from which archaeobotanists can infer hallmarks of domestication such as loss of spike brittleness^3^. Molecular geneticists have isolated the genes that control this and other domestication traits and studied sequence diversity at the surrounding loci in wild and domesticated forms to pinpoint probable wild source populations^4^. Genome sequences are an important resource for crop evolutionary research^5^. As whole-genome sequencing has become more affordable in the past two decades, the field has blossomed. Many crop evolutionists operate, often tacitly, under the assumption that present-day population structure, especially in wild relatives, can reflect the situation thousands of years ago. No such proviso is needed for methods such as the pairwise sequentially Markovian coalescent (PMSC) that infer historic population size trajectories from genome sequences of extant individuals^6^. A recent addition to the population geneticist’s toolkit is IntroBlocker, a software that defines ancestral haplotype groups (AHGs)^7^. These co-inherited blocks of single nucleotide polymorphisms (i.e. haplotypes) make it possible to assign “closest wild relatives” not only at the whole-plant or population level, but also for single megabase-sized genomic regions. Lastly, ancient DNA sequences can offer a window into past genetic diversity^8^. DNA sequences extracted from well-preserved ancient specimens, mainly bones have repeatedly upended our understanding of human prehistory^9^. Despite some successes^10,11^, ancient plant DNA has had a comparatively lower impact on crop evolutionary studies owing to the poor preservation of plant material in most climates^12^.

Where and when plants were first domesticated are questions of great importance. In the past, the quest for that single point in time and space when domestication took place has fueled geneticists’ efforts to map “centers of origin” with molecular markers. Such attempts in Einkorn wheat^13^ were met with conceptual and methodological criticism^14^. The point is moot in barley. Several lines of evidence rule out a monophyletic origin of this crop. One is the presence of two independently acting loss-of-function mutations that abolish spike brittleness. The closest wild relatives of these causal mutant alleles are found today in geographically and genetically separated wild populations^4^. Further evidence comes from genome-wide marker data that led Poets et al.^15^ to propose a model of mosaic ancestry, in which domesticated barley is not descended from a single source but has received contributions from different wild populations. Poets et al.^15^ and others^16,17^ studied barley evolution before mostly complete and highly contiguous sequence assemblies of the barley genome^18^ became available and relied on markers ascertained in domesticated barley or reduced presentation sequencing. Here, we use whole-genome sequences of diverse wild and domesticated barleys, including ancient specimens, to connect haplotypes in wild and domesticated populations to answer questions such as: Which regions of the genome trace back to which wild ancestors? Which wild barley genomes harbor the closest present-day wild relatives of domesticated haplotypes? How have haplotypes been reshufled after domestication?

## Structure and divergence of wild barley populations

We started with the assumption that the present-day population structure of wild barley is related to what it was when human beings began to grow barley. Wild barley (*Hordeum vulgare* subsp. *spontaneum*) is a genetically diverse taxon that occurs throughout Western Asia. We sequenced a total of 380 wild barley accessions, many of them from the Wild Barley Diversity Collection, to 10-fold coverage with Illumina short reads (**Supplementary Table 1, 2**). Previous studies on wild barley agree on the fact that isolation-by-distance is the main driver of population differentiation in wild barley^17,19^. Using model-based ancestry estimation^20^ complemented by principal component analysis^21^ (PCA), we divided our panel into five populations whose geographic distributions roughly trace a path from the Southern Levant (SL), via the Syrian Desert (SD), the Northern Levant (NL), Northern Mesopotamia (NM) and Central Asia (CA) (**Fig. 1a, Extended Data Fig. 1a-c, Supplementary Table 3**). These populations had different levels of diversity (**Extended Data Fig. 1d, Supplementary Table 4)**. Low diversity in the SD populations, which was accompanied by high differentiation from other populations, might be explained by higher genetic drift in SD **(Extended Data Fig. 1e)**.

**Figure 1:**
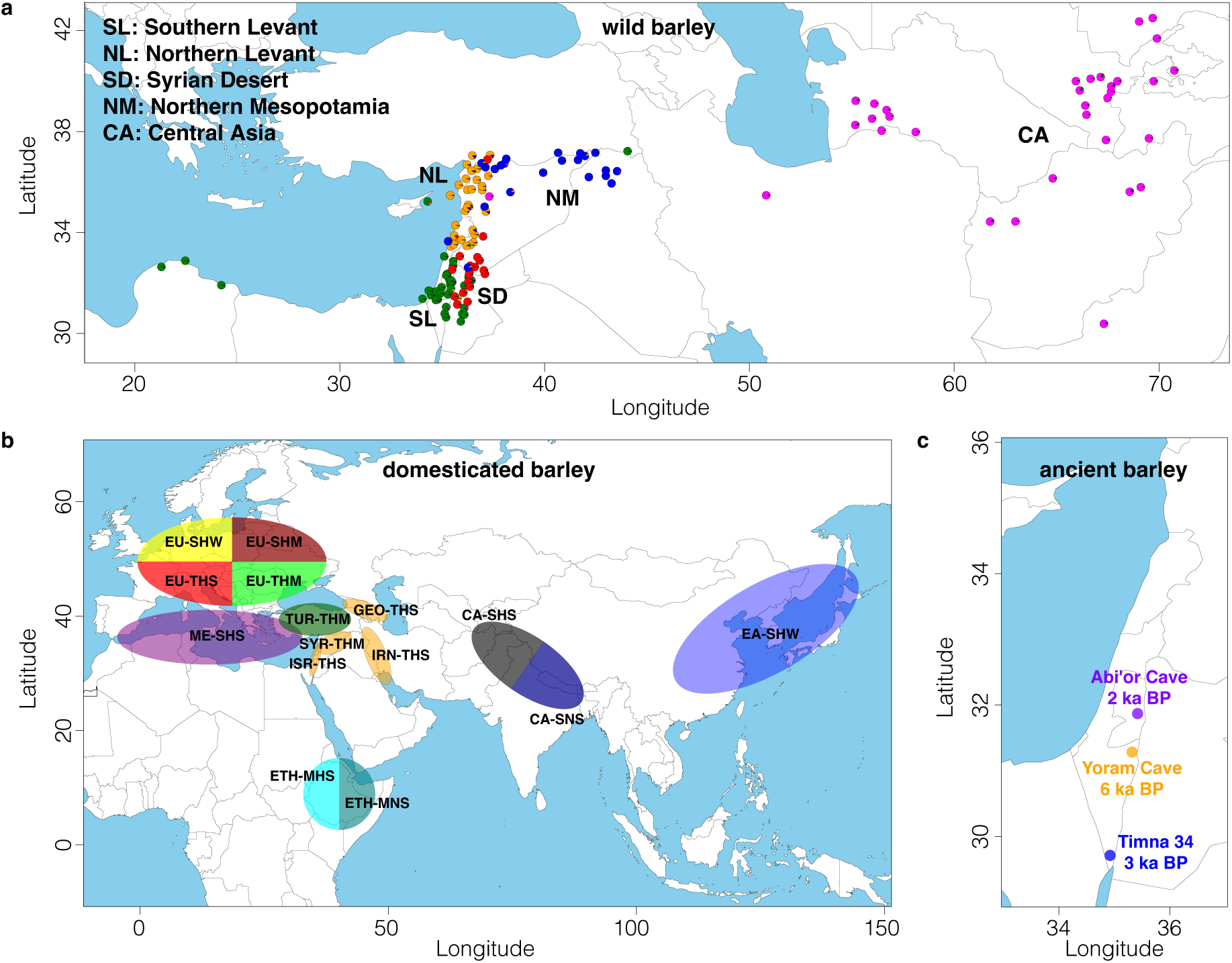
Diversity panel of wild and domesticated barley. **(a)** Collection sites and population structure of 143 wild barley genotypes with precise geographical locations. Pie charts show the results of model-based ancestry estimation with ADMIXTURE (K=5) and are plotted at approximate collection sites. Jitter was added to avoid overlaps between nearby accessions. Only unadmixed samples, i.e. those whose major ancestry components was ≥ 0.85 are shown. **(b)** Assignment to macrogeographic regions of 15 populations inferred from GBS data of 19,778 domesticated barley^26^. The population names encode the samples’ most common origin and their predominant morphological and phenological characters (row type, lemma adherence, annual growth habit) as detailed in **Supplementary Table 5**). **(c)** Archaeological sites in the Judean Desert at which ancient barley grains used for ancient DNA extractions were found. Ages of the samples, as determined by radiocarbon dating, are indicated in the figure.

If there were no recombination and gene flow, the number of sequence variants between two genomes would inform directly about divergence times. Three examples illustrate that this simple model is not applicable in barley: when we counted single-nucleotide polymorphisms (SNPs) in 1 Mb windows and plotted the SNP distribution, we observed, between some pairs of samples, local differences in divergence times, most prominently between distal and proximal regions (**Extended Data Fig. 2a,b**). In barley and its relatives wheat and rye, proximal non-recombining regions, so-called “genetic centromeres”, are extensive, have fewer genes and drastically reduced recombination^22–24^. In domesticated barley, sequence diversity in these regions is lower, too^17,22^. The situation in wild barley is more nuanced. Looking only at between-population comparisons, the distributions of divergence times were unimodal in distal regions of all chromosomes with a peak at around 600 thousand years before the present (ka BP), which corresponds to a trough in effective population size at the same period (**Fig. 2a,b, Extended Data Fig. 2c**). Fluctuations of population size were also evident from historic trajectories of effective population sizes computed with PSMC^6^ (**Fig. 2b, Extended Data Fig. 3b**). These data indicate that all wild barley populations have recovered from a bottleneck between 2000 and 500 ka BP. A later bottleneck (110 to 120 ka BP) coincided with the Last Glacial Maximum (LGM).

**Figure 2:**
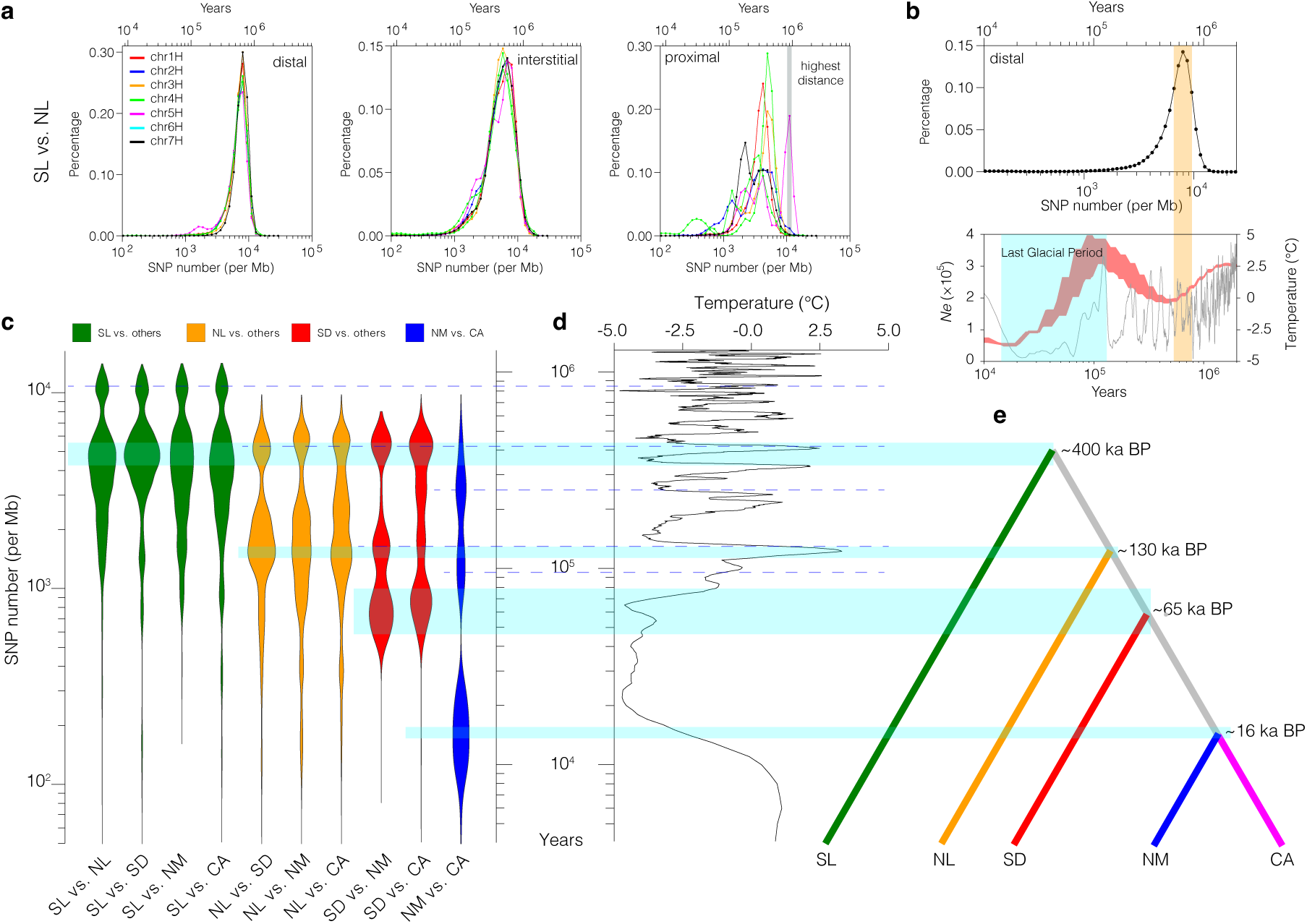
Evolutionary history of wild barley. **(a)** Distribution of sequence divergence (SNPs per Mb) between pairs of accessions from the SL and NL populations in distal, interstitial and proximal regions. The grey shading in the right-hand panel marks the highest divergence between both populations owing to the presence of deeply diverged haplotypes on chromosome 5H. **(b)** The top panel shows the distribution of pairwise sequence divergence for all sample pairs in distal regions of the genome. The bottom panel shows the historic trajectories of effective population sizes in wild barley as inferred by PSMC (red) and global average surface temperatures^39^ (gray). The orange shading marks a simultaneous decline of population size and temperature that corresponds to peaks in the SNP distribution. The last glacial period (100 ka BP to 12 ka BP) is marked by blue shading. **(c)** Violin plots showing the distributions of pairwise sequence divergence in proximal regions of five wild barley populations. Blue shading highlights the peaks in the distribution that mark the most recent divergence between pairs of populations. Earlier such events are marked by dashed lines. **(d)** Global average surface temperatures^39^ in the past 2 million years. **(e)** The divergence of barley populations (most recent inferred split times) is represented as a tree.

Distributions of divergence time in proximal regions were multimodal and differed between chromosomes (**Extended Data Fig. 2c**). This observation defies easy explanation. It may stem from the paucity of centromeric haplotypes and their persistence as single linkage blocks on evolutionary time scales. We took advantage of this rather peculiar situation and delved into the relationship between the divergence of long centromeric haplotypes and that of between individuals and split times between populations (**Supplementary Fig. 1).** To do so, we used SNPs in pericentromeric regions (centromere +/- 25 Mb) to calculate pairwise divergence times between wild barley individuals and arranged wild barley populations in a tree structure based on their most recent splits from each other (**Fig. 2c,e, Extended Data Fig. 3a**). This is a simplification of the relationships between barley populations because divergence times were multimodal and the peaks of the distribution aligned with fluctuations in global surface temperature (**Fig. 2d**). This pattern may be attributable to repeated episodes of colonization of new habitats, contraction and potential loss of populations, recolonization and secondary contact between populations. For example, the common ancestor of the Syrian Desert, Northern Mesopotamian and Central Asian populations split from the Northern Levantine lineage around 120 ka BP when a warm climate may have created new habitats. The Northern Mesopotamian and Central Asian populations split around 17 ka BP. This is consistent with the paleoclimatic modelling of Jakob et al.^19^, according to whom wild barley was absent from Central Asia as recently as 21 ka BP. The old age of the Southern Levantine population (i.e. its early divergence from populations elsewhere) is consistent with that region’s supposed status as a glacial refugium^19^. We were intrigued by the presence of a centromeric haplotype in some Southern Levantine wild barleys that diverged from other such haplotypes around 900 ka BP (**Fig. 2a, Extended Data Figs. 2c, 4**). This is much a deeper split than seen within and between other wild barley populations. The ‘relict’ haplotype may be a chance escape from genetic drift owing to larger population sizes in the Southern Levant or may have been retained by selection for some adaptive advantage it confers. The latter hypothesis is lent some support by the fact that the relict haplotype predominates in many domesticated barley populations (**Extended Data Fig. 4d**). Fang et al.^25^ speculated that higher-than-average differentiation between wild barley 5H may have been caused by a large pericentric inversion on that chromosome. We did see inversions in this region, but they did not extend across the entire haplotypes and occurred in other haplotypes (**Extended Data Fig. 4e**), making it unlikely that structural variation is the sole explanation for the long persistence of the relict haplotype.

## A haplotype-based view of barley evolution

To add domesticated barley to the picture, we selected from a large collection of 19,778 domesticated barley accessions^26^ a panel of 302 samples, of which we sequenced 116 to about 10-fold coverage and 186 to about 3-fold whole-genome coverage (**Fig. 1b**, Online Methods, **Supplementary Figs. 2-4, Supplementary Table 1, 2, 5, 6**). We ran IntroBlocker on these data. As was observed in wheat, sequence divergence in domesticated barley, in contrast to its wild relative, was bimodal. This was true irrespective of whether distal or proximal regions were considered (**Fig. 3a, Extended Data Fig. 5a**). The recent peak at around 98 SNPs per Mb (∼8000 years of divergence) corresponds to a bottleneck that marks the coalescence of many haplotypes into common ancestors in the hypothetical domesticated founder population(s). The earlier peak (6,500 SNPs per Mb, 530,000 years) mirrors that seen in wild barley and arises from comparisons between haplotypes that diverged before domestication. To group haplotypes according to whether they split before or after domestication, we set a threshold of 400 SNPs per 1-Mb window (corresponding to a divergence time of 32,000 years, **Fig. 3a**). We give exemplary figures drawn with a 5 Mb window size (**Fig. 3b, Extended Data Fig. 5b**), but used 100 kb windows after inspecting haplotype length around a key domestication gene (**Supplementary Fig. 5, Supplementary Table 7**).

**Figure 3:**
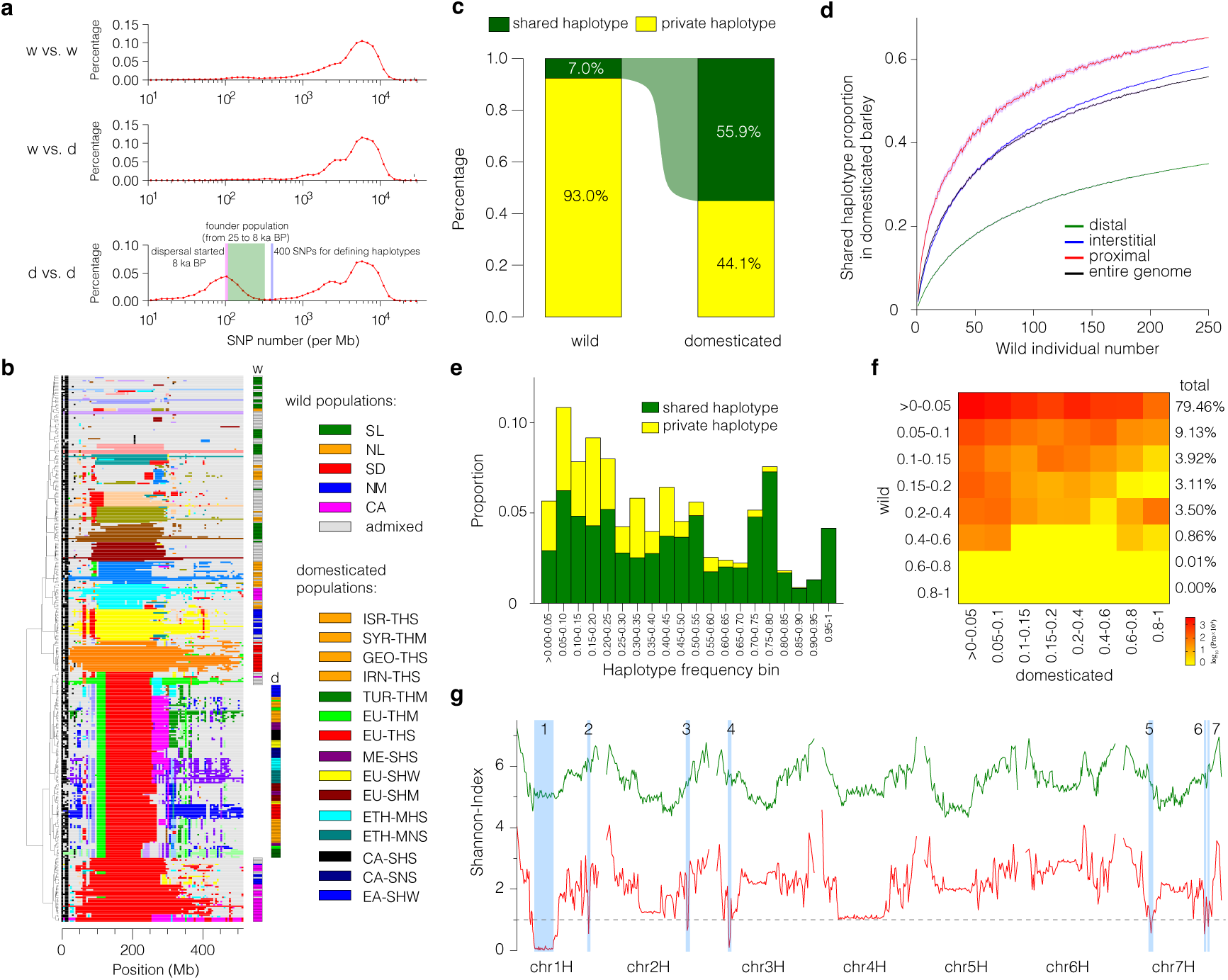
Haplotype diversity in wild and domesticated barley. **(a)** Distribution of sequence divergence (SNPs per Mb) between pairs of wild (w) and domesticated (d) barley. The blue line marks the threshold (400 SNPs per Mb) to delineate the post-and pre-domestication origin of haplotypes in IntroBlocker. The green shading marks the persistence of a hypothetical founder population of diverse wild ancestry that split up into geographically isolated populations from 8 ka BP onwards (purple line). **(b)** Ancestral haplotype groups (AHGs) on chromosome 1H as inferred by IntroBlocker using 5 Mb non-overlapping windows. The 20 most frequent AHGs are shown in different colors. Gray color is used for less frequent AHGs. Colored bars on the right-hand side assign samples to wild (w) or domesticated (d) subpopulations. **(c)** Proportions of shared and private haplotypes in wild and domesticated barley. **(d)** Saturation curves show how the proportion of shared haplotypes in different genomic compartments increases with the number of potential wild barley counterparts. Solid lines and shading denote the average and 95% confidence intervals of 100 random samples. **(e)** Proportions of haplotypes of domesticated barleys that are shared with wild barley at different frequency bins. **(f)** Normalized two-dimensional haplotype frequency spectrum in wild and domesticated barley. The value (percentage) in each cell was determined by dividing counts by the total number of shared haplotypes. Percentages at the right margins indicate the relative sizes of frequency bins in domesticated barley (row sums), e.g. 79.46% of shared haplotypes occurs at 5% frequency or less in wild barley **(g)** Haplotype-based Shannon indices in wild and domesticated barleys. Seven regions with values below 1 in domesticated samples (dashed line) were defined as putative selective sweeps. The genome sequence of B1K-04-12 was used as a reference for this analysis.

A prominent feature in the whole-genome AHG maps of barley was the presence of long centromeric haplotypes that were shared between wild and domesticated barley. This haplotype sharing lends immediate visual support to the notion of the mosaic ancestry of domesticated barley (**Fig. 3b, Extended Data Fig. 5b**). Owing to the lower diversity of haplotypes in proximal than in distal regions of the genome in both wild and domesticated barley, our diversity panel covers nearly all pericentromeric haplotypes, but does not achieve saturation in distal regions (**Extended Data Fig. 6a**). For example, there was only a single pericentromeric haplotype in domesticated barley on chromosome 1H, which was found mainly in Central Asian wild barleys (**Fig. 3b**). To paint a more general picture, 55.9% of domesticated haplotypes were present in at least one wild barley sample; in the converse scenario, 7.0% of wild haplotypes were shared with a domesticated barley (**Fig. 3c**). A saturation analysis makes it seem likely that a larger sample of wild genotypes might unearth more shared haplotypes (**Fig. 3d**). On the other hand, some domesticated haplotypes may lack a wild counterpart: haplotypes private to the domesticate are most common in distal regions of the chromosomes and tend to be rare (**Fig. 3e**). They may have arisen after domestication by recombination of haplotypes inherited from the wild progenitors or their progenitors may have been extinct in the wild because of genetic drift. As expected after a bottleneck, the haplotype frequency spectrum differs between wild and domesticated barley. Common haplotypes (i.e. those with a major allele frequency above 20%) are seldom seen in wild barley, but were more frequent in the domesticate (**Extended Data Fig. 6b**). Still, 79% of haplotypes in domesticated barley with an identifiable wild counterpart occur at low frequency (< 5%) in the wild (**Fig. 3f**). Seven regions of the genome showed an extreme reduction of haplotype diversity in domesticated but not in wild barley (**Fig. 3g**). We inspected local haplotype structure (**Supplementary Fig. 6**) and annotated the functional effects of genomic variants residing in these intervals to prioritize genes for future inquiry (**Supplementary Table 8)**, even though the large sizes of the regions preclude the confident identification of any single plausible candidate gene. More generally, the high genetic differentiation, evident at the level of both SNPs and haplotypes (**Extended Data Fig. 8c, d**), may make it impossible to map selection sweeps by outlier scans: in pairwise comparisons between domesticated populations, on average 7.5% of the genome did not share any haplotypes (**Supplementary Table 9**). Rather than from pervasive forces of adaptive evolution, we suspected that local lineage sorting may underlie this pattern.

## The origins of domesticated haplotypes in time and space

We enquired into the temporal and spatial origin of haplotypes in domesticated barley by running IntroBlocker with different thresholds corresponding to divergence time brackets and inspecting which extant wild barley genomes harbor the closest relatives of domesticated barley (**Fig. 4, Extended Data Fig. 7**). The resultant genome map of spatiotemporal relations is again testimony to the mosaic genomic constitution of the crop (**Fig. 4a, Extended Data Fig. 7**). This pattern established itself early in the evolution of cultivated barley. About 91% of domesticated haplotypes with a wild counterpart split from the latter between 32 and 8 ka BP, i.e. during the formation of the immediate wild progenitor of domesticated barley and the initial stages of domestication (**Fig. 4b**). Fewer than 9% are attributable to more recent gene flow. All five wild barley populations contributed to domesticated barley, albeit in different proportions. Wild barley populations from the Southern and Northern Levant and Central Asia each contributed between 20 and 27 % of haplotypes and those from the Syrian Desert and Western Asia contributed 16.3% and 12.9%, respectively (**Fig. 4b**). There were also differences between domesticated barleys as to how much certain wild populations contributed genetic material to them. Haplotypes from Central Asian wild barleys were found more frequently in domesticated barleys from East and Central Asia than in other domesticated populations (**Fig. 4d**). This close affinity between wild and domesticated barley from “the East” had been noted by Morrell et al.^27^, who saw it as evidence for a second centre of barley domestication east of the Zagros mountains in Iran. Our explanation is that this trend occurred due to gene flow from local wild populations into already domesticated populations coming from the Western Fertile Crescent. The Northern Levantine wild barley population contributed more to domesticated forms in Western Asia and Europe than to those in East and Central Asia, which had more Central Asian ancestry. Mediterranean barleys had a higher share of Southern Levantine ancestry. This relationship may suggest different points of departure of early farmers from the Fertile Crescent. These results are qualitatively similar to those of Poets et al.^15^, but differ in that their analysis, based on 5,000 SNP markers, assigned a greater contribution (> 50%) of Southern Levantine wild barley to all domesticated populations.

**Figure 4:**
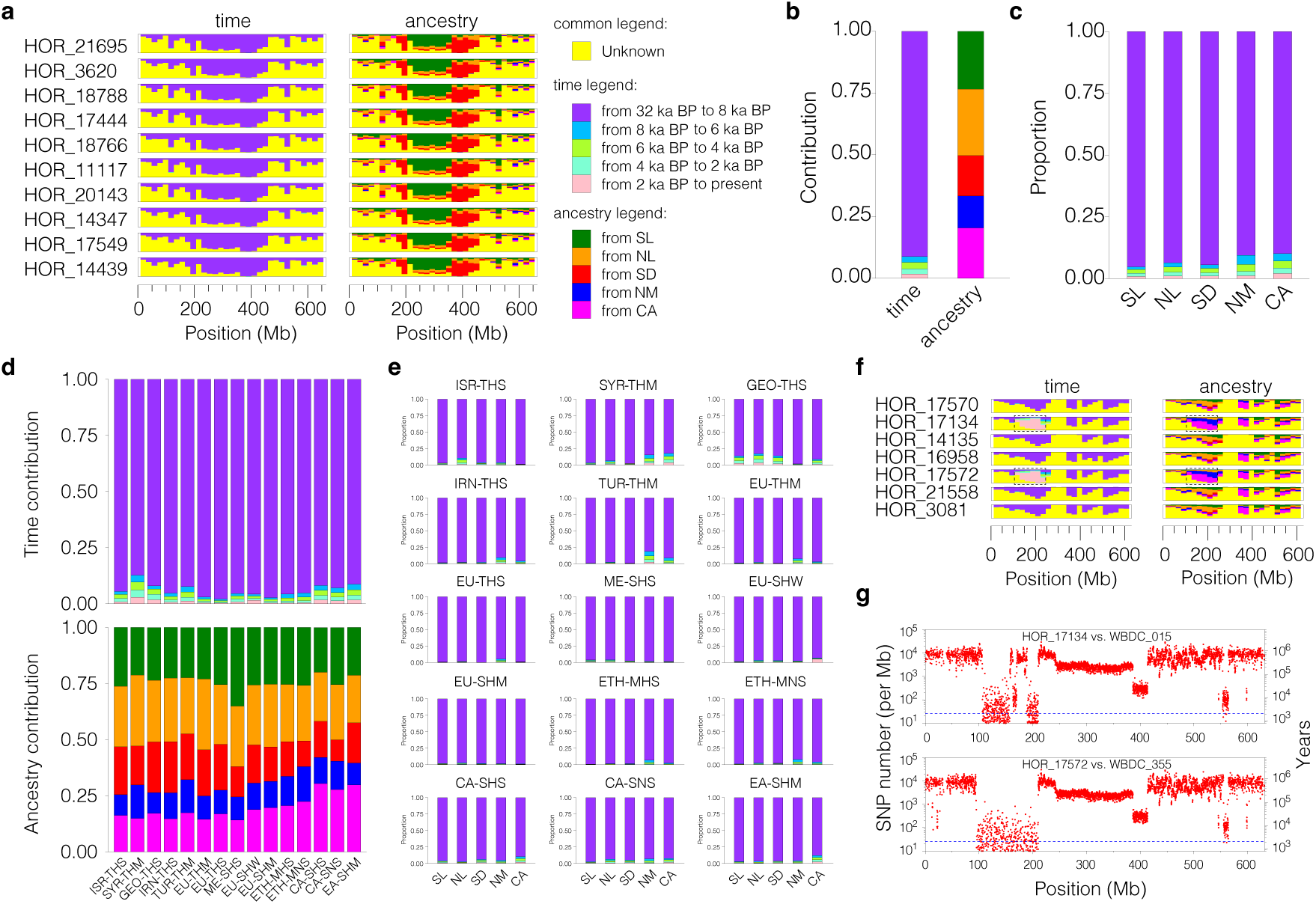
Spatial and temporal origins of haplotypes in domesticated barley. **(a)** Spatiotemporal origins of haplotypes of the EU-THS (European two-rowed spring barley) population of domesticated barley on chromosome 2H in 20-Mb windows. The height of the colored bars is proportional to the probability that a haplotype entered the domesticated gene pool in a certain time period (left panel) or from a certain wild barley population (right panel). Yellow denotes haplotypes of unknown provenance owing to missing data, lack of a clear wild counterpart or potential geneflow from domesticated to wild barley. The results for all domesticated populations are shown in **Extended Data Figure 7. (b)** Spatiotemporal origins of domesticated barley haplotypes across the entire genome. **(c)** Time periods at which haplotypes from each of the five populations entered the domesticated gene pool. **(d)** Spatiotemporal origins of haplotypes in 15 domesticated barley populations. **(e)** Time periods at which haplotypes from each of the five populations entered 15 domesticated barley populations. Haplotypes of unknown provenance were ignored when considering proportions in panels **(b)** to **(e)**. **(f)** Spatiotemporal origins of haplotypes of the EU-SHW (European six-rowed winter barley) population on chromosome 7H. The dashed rectangle marks a haplotype that owes its presence in domesticated barley to recent gene flow. **(g)** Sequence divergence (SNPs per Mb) on chromosome 7H between two EU-SHW accessions (HOR 17134 and HOR 17572) and two Central Asian wild barleys (WBDC 055 and WBDC 355). The dashed line marks 2 ka BP of divergence (random mutation rate: 6.13 SNPs per Gb per generation).

Domesticated barleys differ also in how much recent gene flow they have received from wild barley (**Fig. 4c, e**). Wild introgressions are most common in cultivated accessions from Western and Central Asia, where wild barley is common: 12.8% of haplotypes in Syrian barleys (SYR-THM) are attributable to recent (later than 8 ka BP) wild introgressions (mainly from the Central Asian and Northern Mesopotamian populations). We were surprised to see wild haplotypes flowing into Northern European barley in apparently recent times: the cultivar ‘Kiruna’ (HOR 17134) shared a haplotype on chromosome 7H, 100 – 200 Mb with a Central Asian wild barley (**Fig. 4f, g, Supplementary Fig. 7**). This observation can be explained by the use of wild barley as a genetic resource by breeders: ‘Kiruna’s’ pedigree features ‘Vogelsanger Gold’, a variety from the 1960s with a wild barley introgression^28^. The same haplotype is seen in HOR 17572, which is purported to be an Austrian landrace. We consider errors in the passport records or accidental outcrossing during *ex situ* management the most likely explanation for this case.

## Relationships between domesticated lineages

We inspected divergence levels between haplotypes post-domestication to infer split times between different populations of domesticated barley in a hierarchical manner (**Fig. 5a,b, Extended Fig. 8a, b, Supplementary Table 10**). We used only SNPs in haplotypes descended from the same wild lineage to compute pairwise divergence times between samples. First, we divided our domesticated barley panel into three groups: Western (Near East + Europe), Eastern and Ethiopian barleys, which all diverged from each other around 8.5 ka BP, reflecting the dispersal of agriculture from the Fertile Crescent around that time. Subsequently, Western barley split into three lineages (Near East, two-rowed Europe and six-rowed Europe) around 7.5 ka BP. This is consistent with the archaeological records that show that by 7 ka BP barley had been introduced to Europe, North Africa and Central Asia^29^. These populations subdivided further between 7 and 5 ka BP. Divergence time distributions had multiple peaks in some comparisons. In the case of European barleys, gene flow between populations, which are differentiated by morphology and phenology rather than by geography, is plausible. In the case of Western Asian populations from Georgia and Iran (GEO-THS, IRN-THS) fine-scale population is conceivable: landraces in these mountainous regions may trace back to a common source population but have evolved in mutual reproductive isolation after reaching their current habitats. In **Fig. 5c**., we give a graphical summary of these results in relation to known dispersal routes supported by archaeological evidence.

**Figure 5:**
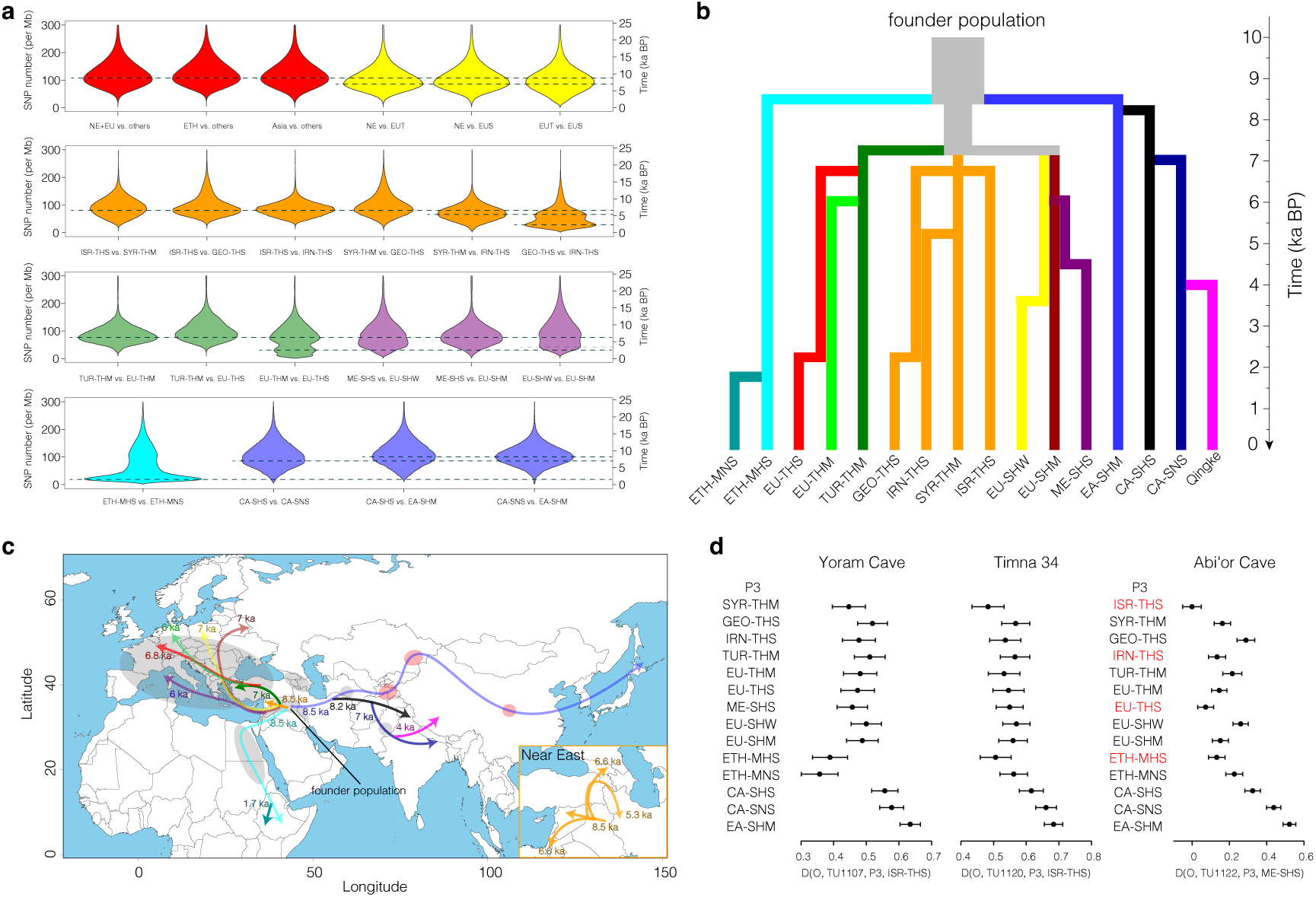
Divergence and dispersal of domesticated barley. **(a)** Violin plots showing the distribution of sequence divergence (SNPs per Mb) in pairwise comparisons between samples from different populations of domesticated barley. Dashed lines mark the peaks of the distributions (split times). Multimodal distributions may have risen from episodes of gene flow. **(b)** Schematic diagram illustrating the lineal descent and split times between 15 barley populations defined in this study and Tibetan barleys (Qingke) studied by Zeng et al.^40^. **(c)** Map showing when and along which routes domesticated barley spread from its center of origin in the Fertile Crescent. Gray shading indicates barley archaeological sites dating back about 7,000 years; red shading indicates barley archaeological sites dating back about 5,000 years^29^. **(d)** D statistics for different comparisons among ancient barleys and 15 domesticated barley populations. The outgroup (O) was *H. pubiflorum*. For samples from Yoram cave and Timna, P_4_ was ISR-THS population; for that from Abi’or cave it was ME-SHS. Positive values indicate that the ancient samples shared more derived alleles with P4 than with P3. Red color marks comparisons that were not statistically significant (Z score below 3).

## A single-gene view of mosaic ancestry

How we think about barley crop evolution owes much to the genetic dissection of loci at which mutant alleles confer traits that are seen only in the domesticate, namely non-shattering (“non-brittle”) spikes, fertile lateral grains (“six-rowed” spikes) and the loss of lemma adherence to the mature grain (“naked” or “hulless” barley). The corresponding genetic loci are *BRITTLE RACHIS 1 and 2* (ref.)^4^, *SIX ROWED SPIKE 1* (ref. ^30^) and *NUDUM*^31^ with mutant alleles *btr1*, *btr2*, *vrs1.a1 to vrs1.a4* and *nud*. These genes were not identified in genome-wide scans for regions with extraordinarily low haplotype diversity (**Extended Data Fig. 6c,d**). The reason for this is that multiple independent loss-of-function of alleles are present at the *BTR1/2* and *VRS1* loci and that the widespread cultivation of naked barleys is confined to a few geographic regions such the Himalayas and Ethiopian highland. Even so, the persistence of long haplotypes (**Extended Data Fig. 9a)** around these genes and the accumulation in them of rare variants since the most recent, and indeed recent (< 10 ka), common ancestor allowed us to date, in an approximate manner, the origin of domesticated loss-of-function alleles. To infer the approximate age of these haplotypes, we inspected distributions of pairwise SNP numbers and translated them into divergence times (**Extended Data Fig. 9a**).

Our estimated age of 25 ka BP for the *btr1* haplotype (**Extended Data Fig. 9a**) predates the earliest archaeobotanical remains of domesticated barley by some 15,000 years^2^. It is not impossible that non-shattering barleys (and the causal haplotypes) languished as rare variants in the wild before early cultivators selected them for propagation. Even though the precision of molecular dating is limited by uncertainties surrounding mutation rate estimates^32^, we can propose the following relative order of emergence of mutant alleles and their surrounding haplotypes: *btr1, btr2, vrs1.a1, nud, vrs1.a3, vrs1.a2,* and *vrs1.a4* (**Extended Data Fig. 9a**). Their most closely related wild counterparts (**Extended Data Fig. 9b, c**) were found in different present-day wild barley populations: Southern Levant (*btr1*, *nud, vrs1.a3*), Northern Levant (*btr2*, *vrs1.a2*) and Northern Mesopotamia and Central Asia (*vrs1.a1*, *vrs1.a4*). This result aligns with earlier gene-based analyses of the *btr1*/*btr2* locus by Pourkheirandish et al.^4^, who posited two origins of tough-rachis barleys, one in the Northern and the other in the Southern Levant. The early origin of the *nud* mutation is consistent with the fact that hulless barleys from places as far apart as Tibet and Ethiopia all share the same 17-kb deletion spanning the *NUD* genes (**Extended Data Fig. 9a**). Yet their overall genomic composition is quite different: the ETH-MNS and CA-SNS population do not share any haplotypes in 44.8% of the genome. We speculate that before the respective ancestors of Central Asian and Ethiopian barleys left the Fertile Crescent, they acquired the common *nud* allele as it was spreading from a single Southern Levantine source across barley’s early gene pool.

## Persistent population structure revealed by ancient DNA

We analyzed ancient DNA sequences of 23 barley grains from the Southern Levant (**Fig. 1c, Supplementary Table 11**) dated to between 6000 and 2000 calibrated years before present (cal BP) to see how they might complement our haplotype map of extant genomes. The short lengths of the sequenced fragments, nucleotide misincorporation profiles and good mapping rates (**Supplementary Table 11, Supplementary Fig. 8**) of the ancient DNA reads to the barley reference genome sequence confirmed their authenticity. All ancient barleys grouped together with cultivated types in a PCA (**Extended Data Fig. 10a**) and had the domesticated *btr1Btr2* haplotype, common in Western barleys (**Supplementary Table 11**). The barleys from Yoram Cave (ca. 6 kya) and Timna Valley Site 34 (ca. 3 kya) were two-rowed forms with the *Vrs1.b2* allele, likewise common in Western types (**Supplementary Table 11**). Those from Abi’or cave (ca. 2 kya) carried the six-rowed (*vrs1.a1*) allele. We used IBS with GBS data (**Supplementary Table 12**), ADMIXTURE^20^ and D-statistics^33^ (**Supplementary Table 13**) to understand the relationship between our ancient samples and present-day barley populations (**Fig. 5d, Extended Data Fig. 10b,c**). The ancient two-rowed barleys were most closely related to extant Western Asian populations, whereas the six-rowed ones were genetically similar to Mediterranean barleys, but also showed Western Asian and European ancestry components (**Extended Data Fig. 10c**). Since the grains from Abi’or cave were dated to 2000 calibrated years BP, i.e. to the Roman period, secondary contact between geographically distant barley population may have been mediated by sea-borne trade across the Mediterranean. Our ancient samples come from a narrow geographic region and do not inform about Eastern or Ethiopian barley. Yet still, the data show that two-and six-rowed Western barleys were genetically distinct already thousands of years ago and that diversity at that time can be meaningfully interpreted in the context of modern barley diversity. All ancient barleys except TU1120 had the same pericentromic haplotype on chromosome 1H as extant domesticated barley. This haplotype is most likely of Central Asian origin (**Extended Data Fig. 7**) and is thus likely to have been introgressed into Southern Levantine barley through gene flow and recombination in a hypothetical founder population. This observation confirms earlier findings^11^ that 6,000 year-old barley resembled present-day domesticated barley more than it did local wild barley populations.

## Discussion

Pankin et al^34^. proposed two models for how the mosaic ancestry of domesticated barley may have come about: (i) recurrent introgressions from different wild populations into an early domesticated (“proto-*vulgare*”) lineage, or (ii) population structure in the wild population(s) from which the domesticated gene pool is derived. As noted by Pankin et al., these models are not mutually exclusive and our analyses support that a combination of the two may have been at play. The geographically diverse origins of the haplotypes surrounding key domestication genes (*btr1/2*, *vrs1*, *nud*) are consistent with (i) a poly-centric initial phase of domestication and (ii) a protracted phase during which barley cultivation remained confined to the Fertile Crescent and local barley populations (along with human farming populations) established themselves throughout the region (**Fig. 6a**). However, this estimated early age of the non-shattering *btr1* haplotype agree with a high frequency (36 %) domestic-type abscission scars in wild barley rachises in the Ohalo II archaeological site (23 ka BP) in the Southern Levant^35^. This is line with the idea that barley cultivation may have begun before the fixation of hallmark domestication traits such as non-brittle spikes^35,36^. A protracted “proto-domestication” is also consistent with a wide trough in trajectories of effective population sizes between 25 and 10 ka BP (**Fig 6b**).

**Figure 6:**
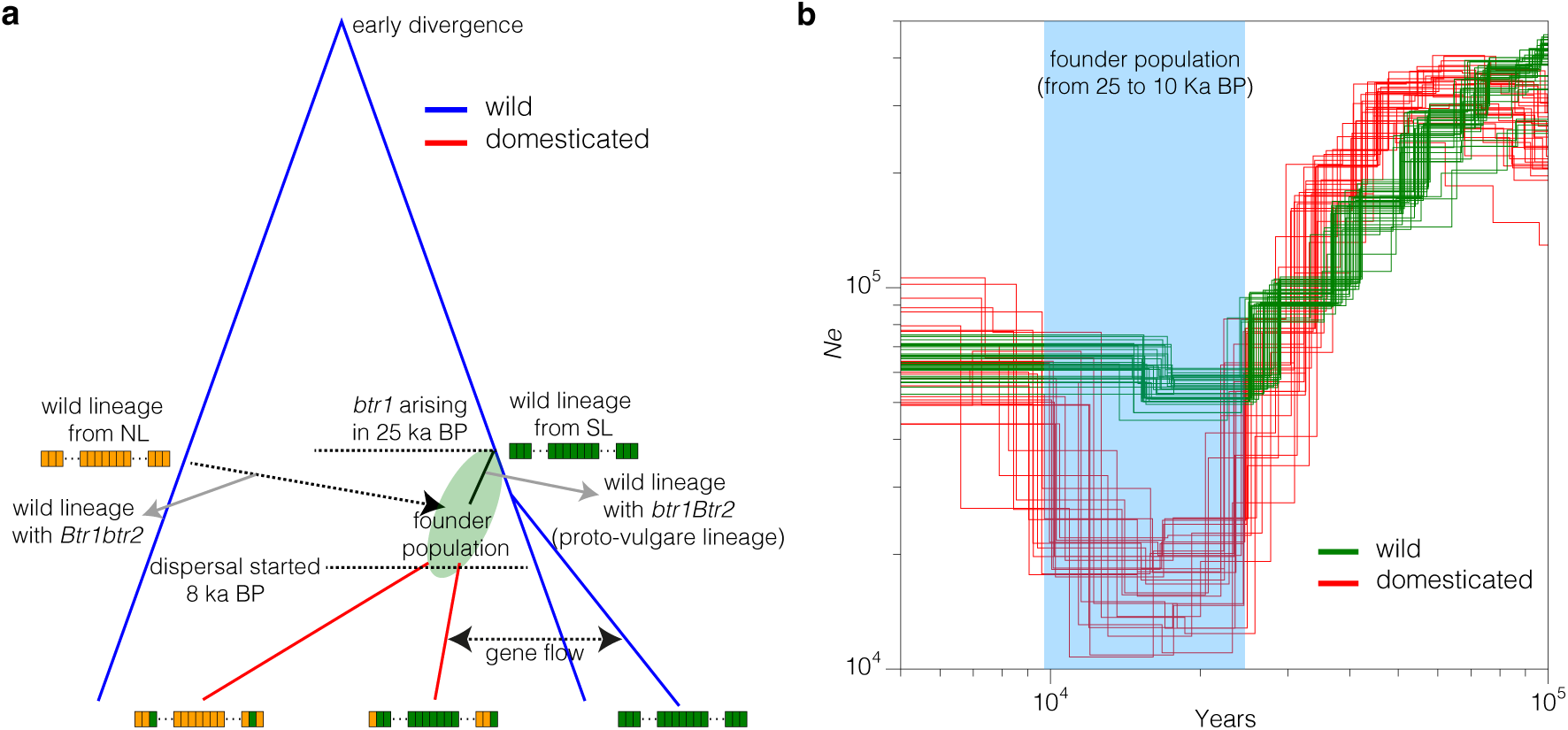
Origin and evolution of domesticated barley. **(a)** Model explaining the origin of the bimodal distribution of sequence divergence in domesticated barley. Two wild populations (SL and NL) are shown for illustration. In fact, all five wild barley populations contributed to the formation of the founder population. Blue and red lines stand for domesticated and wild lineages, respectively. The rectangles represent a chromosome subdivided into windows. Distal and proximal regions are separated by dashed lines. Green and orange genomic windows in extant barleys (bottom) are derived from progenitor haplotypes in the SL and NL populations, respectively. About 25 ka BP, the *btr1* mutation conferring a tough rachis arose in the SL population. The *btr2* mutation arose ca. 22 ka BP in the NL population. Later, non-brittle barley was taken into cultivation independently by early farmers in the respective regions. Gene flow facilitated by human migration or cultural exchange gave rise to an admixed founder population. As farmers and their crops moved out of the Fertile Crescent, the founder population split into several isolated lineages that retained genomic windows of diverse wild ancestry. Later gene flow from wild to domesticated occurs in regions of sympatry. **(b)** Trajectories of historic effective population size inferred by PSMC using wild barley (green) and domesticated barleys (red). Results for all domesticated populations are shown in **Supplementary Fig. 10**.

Local populations of early cultivated barley were sufficiently connected by gene flow to allow for the spread of beneficial genes of large effect. At the same time, crosses between domesticated barley and local wild forms enriched the former’s gene pool, traces of which are seen in present-day patterns of haplotype sharing. For Western Asian domesticated populations, this process continues to the present day and is evident in haplotype exchanges that must have happened long after domestication. Present-day West Asian barleys are thus unlikely to be direct lineal descendants of the early founder population.

As the early agriculturalists moved further from Fertile Crescent, domesticated barley split into separate lineages that evolved largely in isolation, and the influx of wild haplotypes ceased. As current the population structure attests, geography was not only driving force of population splits: farmer’s practices favored distinct gene pools as was the case for European two-rowed and six-rowed barleys. Mosaic ancestry resulting from these processes may pose a limit to the mapping of adaptive loci. As the proto-*vulgare* haplotypes sorted into local lineages, it is likely that adaptation to new habitats also shaped local patterns of sequence diversity. But resolving the effects of both forces may be difficult: in many regions of the genomes, different lineages do not share haplotypes, a pattern also observed in selective sweeps. A promising avenue to disentangle ancestral haplotype structure from adaptive evolution is mutational genomics. An example is the flowering time regulator *HvCENTRORADIALIS*, which was first mapped as a large-effect mutation in a classical barley mutant. Later on, sequence data added geographic range expansion^37^ and newly arisen structural variation^38^ to the picture.

## Acknowledgments

We are grateful for the technical assistance of Susanne König and Ines Walde and IT support by Thomas Münch and Jens Bauernfeind. We thank Zihao Wang and Weilong Guo for their advice on the use of IntroBlocker, and Xinyi Liu and Mona Schreiber for sharing knowledge on the archaeology of barley domestication and dispersal. Andreas Börner supported the development and maintenance of diversity panels. N.S. and M.M. were supported by grants from the German Ministry of Research and Education (BMBF, 031B0190 and 031B0884). B.J.S is supported by the Lieberman-Okinow Endowment at the University of Minnesota. V.J.S. was supported by the University of Zurich’s University Research Priority Program “Evolution in Action: From Genomes to Ecosystems”. E.W. is supported by the Israel Science Foundation (grants No. 1179/13, 545/23).

## Author contributions

M.M, Y.G. and N.S designed the study. A.H. and N.S. conducted and supervised sequencing.

E.W. performed and supervised archaeobotanical work. E.B.-Y. and U.D. performed archaeological excavations. M.D., M.K., A.H.S. and E.W. performed archaeobotanical analyses. V.S., E.R. and J.K. performed and supervised ancient DNA experiments. T.F. and B.J.S. contributed sequencing data. Y.G. and M.J. analyzed data. Y.G. and M.M. wrote the manuscript. All authors edited the manuscript.

## List of supplementary items

**Supplementary Table 1:** ADMIXTURE results (K=4) for 682 barley accessions.

**Supplementary Table 2:** Sequencing statistics of 602 unadmixed barley accessions.

**Supplementary Table 3:** Passport data and population assignment of 251 high-coverage wild barley samples.

**Supplementary Table 4:** Genetic diversity of five wild barley populations.

**Supplementary Table 5:** Accessions from the “orange” subpopulation of Milner et al.^26^ that we selected for whole-genome sequencing.

**Supplementary Table 6:** Passport data and population assignment of 302 domesticated barley samples.

**Supplementary Table 7:** Haplotype numbers and proportions of shared haplotypes in wild and domesticated barley as function of the window size used in the definition of haplotypes.

**Supplementary Table 8:** Putative candidate genes in seven selective sweep regions.

**Supplementary Table 9:** AFD and F_ST_ between pairs of barley populations.

**Supplementary Table 10:** Uni-and multimodal distribution of sequence divergence in domesticated barley populations.

**Supplementary Table 11:** Collection sites, dating and sequencing statistics of 23 ancient samples.

**Supplementary Table 12:** The 20 most closely related extant samples of 23 ancient samples.

**Supplementary Table 13:** D statistics of 7 high-coverage ancient samples and 15 domesticated barley populations.

**Supplementary Table 14:** Boundaries of the genomic compartments in MorexV3.

**Supplementary Table 15:** Composition of the pseudo-diploid genomes used in the PSMC analysis in wild barley.

**Supplementary Table 16:** Composition of the pseudo-diploid genomes used in the PSMC analysis in domesticated barley.

## Methods

### Sample selection for genome sequencing

#### Wild barley

Our wild barley panel (**Supplementary Tables 1 and 3**) is comprised of 285 accessions from the Wild Barley Diversity Collection (WBDC)^1,2^, a collection of ecogeographically diverse accessions. The whole-genome sequencing of the WBDC collection is described in a companion paper by Sallam et al. (<u>provided as Related Manuscript file to the referees</u>). A further 95 diverse barley accessions, mainly from the panel of Russell et al.^3^, were also included. The latter set of samples had been sequenced to 3-fold coverage by Jayakodi et al^4^. Deeper-coverage (∼10-fold) sequence data were collected in the present study.

#### Domesticated barley

Milner et al. defined 12 populations using model-based ancestry estimation with ADMIXTURE^5^ in a global diversity panel of 19,778 domesticated barley, which had been subjected to genotyping-by-sequencing (GBS)^6^. We used the ADMIXTURE results and GBS single-nucleotide polymorphism (SNP) matrix of Milner et al.^6^ for sample selection. Except for the Near-Eastern (NE) population (colored orange in **Fig. 1a** of Milner et al.^6^), we selected samples according to the following procedure. First, un-admixed samples, i.e. those with an ADMIXTURE ancestry coefficient q ≥ 0.95 were used as input for a principal component analysis (PCA) with smartpca^7^ (version 7.2.1). Then, samples were selected to cover the PCA space evenly (**Supplementary Fig. 2**). Owing to its higher genetic diversity and internal substructure, a more sophisticated procedure was followed for the NE population (**Supplementary Fig. 3, Supplementary Table 5**). First, ADMIXTURE^5^ (version 1.23) was run on 1,078 samples of Milner et al.^6^, where the NE ancestry coefficient q was higher than that of all other populations, with q ranging from 0.25 to 0.98. Prior to running ADMIXTURE, the SNP set was thinned with PLINK^8^ (version 1.9) using the parameters ‘--indep-pairwise 50 10 0.1’. For each value of k (number of ancestral population) from 2 to 6, the output of 15 replicate runs of ADMIXTURE with different random seeds was combined with CLUMPP^9^ (version 1.1.2) and plotted with Distruct^10^ (version 1.1). Individuals with q values ≥ 80% for their main ancestry component were considered unadmixed. The results for k=6 was chosen for further analysis. The genetic separation of the defined populations was confirmed with smartpca^7^ (version 7.2.1). Only those samples of the NE sub-populations that were actually located in the Near East were selected for sequencing. The selected samples were distributed in an equidistant manner in the PCA diversity space. In total, we selected 302 samples from 15 populations (**Supplementary Fig. 4, Supplementary Table 6**). The populations were named according to their geographic origins and three key traits closely connected to global population structure^6^: row type (two-rowed [T], six-rowed [S], mixed [M]); lemma adherence (hulled [H], naked [N]); annual growth habit (winter-sown [W], spring-sown [S], mixed [M]). For example, ISR-THS refers to a population whose members are predominantly two-rowed hulled spring barleys from Israel. For each population, we selected about 20 accessions for WGS sequencing. Among these, seven to ten samples of each population (total: 116, “high-coverage samples”) were sequenced to ∼10-fold coverage. The remaining samples of each population were sequenced to ∼3-fold coverage (total: 186, “low-coverage samples”). Seeds for all selected accessions can be ordered from German Federal ex situ genebank at IPK Gatersleben.

### Plant growth, DNA isolation and Illumina sequencing

Plant cultivation and DNA isolation was essentially as described previously^6^. Illumina Nextera DNA Flex WGS libraries were prepared and sequenced (paired-end: 2 x 151 cycles) on an Illumina NovaSeq 6000 device at IPK Gatersleben according to the manufacturer’s instructions (Illumina Inc., San Diego, CA, USA).

### Read mapping and variant calling

The reads of 682 barley genotypes, of which 380 were wild and 302 domesticated, were mapped to the MorexV3 genome sequence assembly^11^ using Minimap2 (ref. ^12^). BAM files were sorted and deduplicated with novosort (V3.06.05) (https://www.novocraft.com/products/novosort/). Variant calling was done with bcftools^13^ (version 1.15.1) using the command ‘mpileup-a DP,AD-q 20-Q 20--ns 3332’. The resultant “raw” SNP matrix was filtered as follows: (i) only bi-allelic SNP sites were kept; (ii) genotype calls were deemed successful if their read depth was ≥ 2 and ≤ 50; otherwise genotypes were set to missing. SNP sites with fewer than 20% missing calls and fewer than 20% heterozygous calls were used for ADMIXTURE runs (with k ranging from 2 to 4) as described above. At k = 4, wild individuals with ≥ 15% ancestry from domesticated barley were considered admixed. A total of 80 wild admixed samples were excluded from subsequent analyses (**Supplementary Fig. 10, Supplementary Table 1**). Mapping statistics of the remaining 602 accessions are shown in **Supplementary Table 2**. A total of 251 wild barley samples with high coverage (∼10x) without domesticated admixture were used for subsequent population genetic analyses.

We prepared two SNP matrices, SNP1 and SNP2, for downstream analysis. For SNP1, we extracted the data for 367 (251 wild, 116 domesticated) high-coverage samples from the raw SNP matrix. SNP1 was filtered as follows: (i) only bi-allelic SNP sites were kept; (ii) homozygous calls were deemed successful if their read depth was ≥2 and ≤50 and set to missing otherwise; (iii) heterozygous calls were deemed successful if the allelic depth of both alleles was ≥5 and set to missing otherwise. The SNP2 matrix contains variants for 302 domesticated samples and was constructed from another bcftools run using the same parameters as above but with a down-sampled data set, in which the read alignments of the high-coverage samples (n=116) had been thinned so as to achieve a sequence depth comparable to that of the low-coverage samples (**Supplementary Fig. 11**) using SAMtools^13^ (version 1.16.1) with the command “samtools view-s 0.FRAC” (FRAC is the sampling rate). The targeted number of uniquely mapped (Q20), deduplicated mapped reads for the down-sampled high-coverage data was set to a random number between 35M and 52M. Note that the read length was 2×150 bp in all samples. The matrix SNP2 was filtered as follows: (i) only bi-allelic SNP sites were kept; (ii) homozygous calls were considered successful if their read depth was ≥ 2 and ≤ 20 and set to missing otherwise; (iii) all heterozygous calls were set to missing. A flow chart describing the construction of the SNP matrices used in this study is shown in **Supplementary Fig. 12.** In analyses where the use of an outgroup was required, we used WGS data^14^ of *Hordeum pubiflorum*. Read mapping and SNP calling were done as described above with one difference: a VCF file for all sites in the genome, including those identical to the reference genome, was obtained. This VCF files was merged with other VCF files to determine ancestral states.

### SNP-based genetic distances

The number of SNPs between any two high-coverage genotypes were calculated as follows. First, pairwise SNP numbers were determined in genomic windows with PLINK2 version (2.00a3.3LM, ref. ^15^) with the command “plink2--from-bp x--to-bp y--sample-diff counts-only counts-cols=ibs0,ibs1 ids=s1 s2…”, where x and y are the start and end coordinates of a window and “s1 s2…” is a list of sample IDs. Different window sizes were used: 100 kb (shift: 20 kb), 500 kb (shift: 100 kb), 1 Mb (shift: 200 kb), 2 Mb (shift: 400 kb), 5 Mb (shift: 5 Mb). Then, in each window, a normalized distance measure was calculated to account for the fact that owing to differences in the mappability of short reads, the effective coverage differs between genomic windows^16^ (**Supplementary Fig. 13**). Per-bp read depth was determined for each sample and each position of the reference genome with the command “samtools view-q 20-F 3332 | samtools depth”. The effectively covered region of each window was defined as the union of sites with read depths between 2 and 50. For each, pairwise comparison between samples, the effectively covered regions were intersected using a Perl script. The pairwise distance in a genomic window was calculated as (hom + het/2)/cov, where hom and het are numbers of homozygous and heterozygous differences, respectively and cov is the size of the intersection of the effectively covered regions of both samples. Genomic windows were considered only if the latter quantity amounted to half the size of the window; otherwise the distance was set to missing.

### LD decay

The barley genome was split into three compartments (distal, interstitial, proximal) based on recombination rates^17^ (**Supplementary Fig. 14, Supplementary Table 14**). LD decay was calculated for both wild and domesticated barley in each compartment using PopLDdecay^18^ (version 3.42) with the command “-Het 0.99-Miss 0.2-MAF 0.01-MaxDist 500”.

### Evolutionary analyses in wild barley

Variants calls of 251 high-coverage wild barley samples were extracted from the matrix SNP1 (see above). SNPs sites with fewer than 20% missing calls, fewer than 20% heterozygous calls and minor allele frequency (MAF) ≥ 5% were used in population structure analysis. Model-based ancestry estimation was done with ADMIXTURE^5^ (version 1.23). The number of ancestral populations k ranged from 2 to 5. PCA was done with smartpca^7^ (version 7.2.1). Genotype calls of the outgroup sample *H. pubiflorum* were merged with the SNP matrix, and an IBS-based genetic distance matrix was calculated with PLINK^8^ (version 1.9). The distance matrix was used to construct a neighbor-joining (NJ) tree with Fneighbor (http://emboss.toulouse.inra.fr/cgi-bin/emboss/fneighbor), which is part of the EMBOSS package^19^. The resultant tree was visualized with Interactive Tree Of Life (iTOL)^20^. In each of the five wild barley subpopulations, the nucleotide diversity^21^ (π) and Watterson’s estimator^22^ (θ_W_) were calculated from the SNP matrix without MAF filtering using a published Perl script^16^. Pairwise fixation indices (*F*_ST_) between pairs of wild barley populations were calculated in genomic windows (size: 1 Mb, shift: 500 kb) using Hudson’s estimator with the formula given as equation 10 in ref. ^23^ using a published Perl script^16^. Coverage-normalized SNP distances were calculated as described above in 1 Mb genomic windows (shift: 200 kb). Distributions of log_10_-transformed distances in the genomic compartments distal, interstitial and proximal were plotted for each wild barley population in R (ref. ^24^, version 3.5.1). To infer divergence times, only SNPs in a 50-Mb region flanking the centromeres (+/- 25 Mb) were used. SNP distances were converted into divergence times using the formula g=d/2μ, where g is the number of generations, μ is the mutation rate and d is the number of SNPs per bp. We assumed that the generation time in the annual species *Hordeum vulgare* is one year. We used a random mutation rate of 6.13×10^-9^ as had been determined by Wang et al.^18^ in the Pooideae grass *Brachypodium distachyon*. The SNP number distribution was visualized by frequency polygons with logarithmic binning (number of bins: 50, range: 10^1^ to 10^4.5^ [31,622 SNPs]. Demographic inference was done with PSMC^25^ (default parameters) using pseudo-diploid genomes, which were created by combining the BAM files of two homozygous individuals as described previously^26–28^. One haplotype was selected from the SL population, the other from one of the other four wild barley populations. A total of 38 pseudo-diploid genomes (**Supplementary Table 15**) were generated. PSMC was run with default parameters. We used MUMmer^29^ (version 4.0.0) to align eight barley genome assemblies with different haplotypes^30^ on chromosome 5H, 100 Mb to 300 Mb. The minimum alignment identity was 90 and the minimum alignment length was 2000 bp.

### Definition of ancestral haplotype groups

Ancestral haplotype groups (AHG) were defined with IntroBlocker^31^. To determine an appropriate threshold for separating haplotypes, we computed coverage-normalized SNP-based distances in 1 Mb windows (shift: 200 kb) (1) among wild samples, (2) among domesticated samples, and (3) between wild and domesticated samples. In each of the three cases, all possible pairwise combinations of samples were considered. We selected a threshold of 400 SNPs per Mb to separate AHGs. Coverage normalized SNP-distance matrices computed from 367 high-coverage samples were as used as input for IntroBlocker with the “semi-supervised” model, giving precedence to wild over domesticated samples in the labelling of AHGs. IntroBlocker was run with different window sizes: 100 kb (shift: 20 kb), 500 kb (shift: 100 kb), 1 Mb (shift: 200 kb), 2 Mb (shift: 400 kb), 5 Mb (shift: 5 Mb). The results of the 5 Mb run are shown in **Fig. 3b** and **Extended Data Fig. 5b**. After inspection of results (**Supplementary Fig. 5**), the results from 100 kb run were used for downstream analyses.

### Analysis of the AHG matrix

The proportions of shared and private AHGs in wild and domesticated barleys were determined with custom Perl scripts. Saturation curves were calculated as follows. We chose sets of k wild barleys (from a universe of 251 samples) at random, with k ranging from 1 to 250. For each k, the selection was repeated 100 times. For each of the samples, we determined the proportion of haplotypes seen in the domesticate that were shared with that set. Mean values and 95 percent confidence intervals for each k were calculated in R (ref. ^24^, version 3.5.1) with a t-test. Two-dimensional haplotype frequency spectra were calculated with custom Perl scripts. Genomic windows with more than 20 % missing data points were excluded.

To infer the times at which wild haplotypes entered the domesticated gene pool, we ran IntroBlocker with different thresholds for haplotype separation: 400 SNPs (equivalent to an approximate divergent time of 32,000 years ago), 98 SNPs (8,000 years), 73 (6,000 years), 49 SNPs (4,000 years), 24 SNPs (2,000 years). Each window in which domesticated and wild samples shared a haplotype, the temporal and spatial origins of that sharing was determined as illustrated in **Extended Data Fig. 7**. For each domesticated haplotype, we compared the results from IntroBlocker runs with different thresholds (divergence time brackets). The latest bracket in which haplotype sharing between wild and domesticated samples occurred was considered a *terminus post quem* for when a wild haplotype type entered the domesticated genepool. This method is agnostic about the direction of gene flow. To exclude recent introgressions from domesticated to wild barley, we removed windows in which multiple domesticated barley samples and a few wild barleys share haplotypes that diverged within the past 8,000 years. To determine the spatial origin of haplotypes, we averaged the ancestry ADMIXTURE coefficients of all wild individuals in which a given domesticated haplotype occurred (**Supplementary Fig. 15**). If two wild samples that shared a domesticated haplotype were highly similar (pairwise IBS >= 0.95), only one was used for the calculation.

### Haplotype-based genetic diversity and selective sweeps

Saturation curves for the average number of haplotypes in a genomic window as a function of sample size were obtained by randomly selecting k individuals with k ranging from 1 to 115 for domesticated samples and from 1 to 250 for wild samples. For each k, the selection was repeated 100 times. Average haplotype numbers were determined for each subsample. Mean values and 95% confidence intervals were calculated in R (version 3.5.1) using a t-test. Watterson’s estimator^22^ (θ_W_) and the Shannon diversity index^32^ were calculated with a custom Perl script on haplotype matrices including only genomic windows with less than 20 % missing data. The θ_W_ and Shannon index in seven barley chromosomes were plotted with Gnuplot using “smooth bezier”.

We looked for regions of reduced diversity in domesticated relative to wild barley and therein searched for genes that might have been potential targets of selection. In order not to bias the analysis by the use of a domesticated reference genome (that of cultivar Morex), we repeated read mapping, variant calling and haplotype inference with the annotated genome of the wild barley B1K-04-02 (FT11)^30^. Regions with a Shannon index ≤ 1 were considered selective sweeps. The effects of SNPs and indels residing in the genes of those regions were classified with SnpEff^33^ version 4.3t and variants with high allele frequency differentiation were prioritized.

The differentiation between populations of domesticated barley was assessed by computing the absolute allele frequency difference (AFD)^34^. The following comparisons were done: NE+EU vs. ETH, NE+EU vs. Asia, ETH vs. Asia, NE vs. EUT, NE vs. EUS and EUT vs. EUS. In addition, we calculated F_ST_ in genomic windows (size 100 kb, shift: 20 kb) using the same method as in wild barley. AFD was used for haplotypes derived from high-coverage samples; F_ST_ calculations were performed for all samples, including low-coverage ones.

### Demographic history of domesticated barley

Trajectories of effective population size across time were inferred with PSMC^25^ (default parameters) using pseudo-diploid genome sequence from two homozygous barley individuals. A generation time of 1 year and a mutation rate of 6.13×10^-9^ were used. First, 27 pseudo-diploid genomes were constructed by combining the sequence data of pairs of individuals from three groups of domesticated barley samples (**Supplementary Table 16**): Europe and Near East; East and Central Asia; and Ethiopia. Here, the pseudo-diploid genomes derived from any of the two of the three mentioned populations and were used to infer the average demographic history of the entire domesticated barley population. Second, we ran PSMC on 341 pseudo-haploid genomes obtained from all possible permutation of sample pairs from within 15 domesticated populations to reflect the population history of each subpopulation of domesticated barley.

Split times between pairs of domesticated barley populations were determined by inspecting the distributions of SNP numbers between pairs of samples in those windows (size: 1Mb, shift: 200 kb) where a given pair of samples differed by fewer than 300 SNPs (corresponding to a divergence of 24,470 years). We only considered windows with an effective coverage of 90 %, i.e. 900 kb or more had at least 2-fold coverage in both samples. The SNP number distribution was visualized by frequency polygons (linear binning; number of bins: 50; range: 0 to 300. SNP numbers were converted to divergence time using the following formula: time = (SNP number per Mb / 10^6^) / (2 ⨉ 6.13 ⨉ 10^-9^), where the 6.13 ⨉ 10^-9^ was the random mutation rate (μ) of *B. distachyon*.

To infer the origins of three genes, we inspected the SNP distribution in sweep haplotypes around the genes and inferred the emergence times of loss-of-function mutations in cultivated barley (**Supplementary Fig. 16**). A Neighbor-Joining tree for each gene was constructed with SNPs from an interval within their sweep region. For the *btr1/2*, *vrs1*, and *nud* loci, the interval extended from 39.4 to 39.7 Mb on chromosome 3H, from 570.5 to 517.2 Mb on chromosome 3H; And 525.3-525.7 Mb on chromosome 7H, respectively.

### Archaeological excavations

We analyzed ancient DNA sequences of 23 barley grains excavated at three archaeological sites in Israel (**Supplementary Table 11**). This number included published data of 5 barley grains from Yoram Cave^35^. Archaeobotanical procedures were performed as described by Lev-Marom et al. Yoram Cave and Timna Valley Site 34 have been described by Mascher et al.^35^ and Lev-Marom et al. Abi’or Cave is a medium-sized cave located on the eastern slopes of the Judean Desert, above Jericho, approximately 50 meters below sea level, across from the Karantal Monastery. The excavations at the cave were directed by the late H. Eshel in 1986. It is situated above a larger cave known as “The Spies Cave”. The cave contains a main long tunnel, approximately 50 meters long, and has revealed archaeological material dating from the Chalcolithic period to the time of the Bar Kokhba Revolt (2nd century CE). The cave was found to be heavily disturbed by animals, antiquities robbers, and monks who lived in it during the Islamic and more recent periods.

### Ancient DNA sequencing and analysis

All laboratory procedures for sampling, DNA extraction, library preparation, and library indexing were conducted in facilities dedicated to ancient DNA work at the University of Tübingen. Before DNA extraction, all seeds were cut into two parts: one part of each seed (36-6.5 mg) was used for DNA extraction and further processing, the other part (26-3.4 mg) was used for radiocarbon dating at the Klaus-Tschira-Archäometrie-Zentrum, Curt-Engelhorn-Zentrum Archäometrie gGmbH, Mannheim, Germany. DNA extraction was then performed according to a well-established extraction protocol for ancient plant material^35^ and double-stranded dual indexed DNA libraries were produced^36,37^. Six ancient DNA samples (TU697 and JK2281-JK3014) were treated with uracil–DNA–glycosylase (UDG)^38^ before sequencing. Sequencing was done on Illumina devices at IPK Gatersleben, the University of Tübingen and the Max-Planck Institute or the Science of Human History Jena. Paired-end Illumina reads of each sample were merged with leeHom^39^ and mapped to the MorexV3 genome sequence assembly with Minimap2 (ref.^12^). BAM files were sorted and duplicates were marked with Novosort (version 3.06.05) (https://www.novocraft.com/products/novosort/). Nucleotide misincorporation profiles were generated with mapDamage2.0 (ref. ^40^). Variant calling was done with BCFtools^13^ (version 1.15.1) using the command ‘mpileup-a DP,AD-q 20-Q 20--ns 3332’. We omitted the parameter “--variants-only” in “bcftools call” to output genotype in all sites. G->A and C->T were excluded, where the G and C are the alleles in the reference genomes and T and A are the alternative alleles called from the short-read data. The resultant SNP matrix was merged with the three different SNP matrices: SNP1 (367 high-coverage samples), SNP2 (302 domesticated barleys) and a published SNP matrix constructed from GBS data of 19,778 domesticated barleys^6^. The matrices had been filtered for site-level missing rate (< 20 %) before merging. The merged SNP1 matrix was used for PCA with smartPCA^7^ (version 7.2.1) using the parameter “lsqproject: YES”. A neighbor joining tree was constructed using only SNPs in the pericentromeric regions of chromosome 1H (150 to 200 Mb in MorexV3) and using only seven ancient DNA samples with high-coverage. The merged GBS matrix was used to compute an IBS matrix with PLINK^8^ (version 1.9). The merged SNP2 matrix was used for (i) ancestry inference with ADMIXTURE^5^ with k, the number of ancestral populations, ranging from 2 to 11; and (ii) the calculation of D statistics with the qpDstat program of ADMIXTOOLS^41^ (version 3.0).

## Code availability

The shell and Perl script used in this study are available from https://github.com/guoyu-meng/barley-haplotype-script.

## Data availability

The sequence data collected in this study have been deposited at the European Nucleotide Archive^42^ (ENA) under BioProjects PRJEB65046, PRJEB56087 and PRJEB53924. The SNP and indel variant matrix will be available at the European Variation Archive^43^ (EVA) under BioProject PRJEB79752. ENA accession codes for individual genotypes are listed in **Supplementary Table 1**. AHG matrices have been deposited in the Plant Genomics & Phenomics Research Data Repository^44^ under a DOI (temporary link for reviewers: https://doi.ipk-gatersleben.de/DOI/a8515674-5c48-41a6-a662-1cb6b7cfff6c/f2a9c92f-972a-4256-be96-e5f028ecdc77/2/1847940088).

---

**Supplementary Figure 1:**
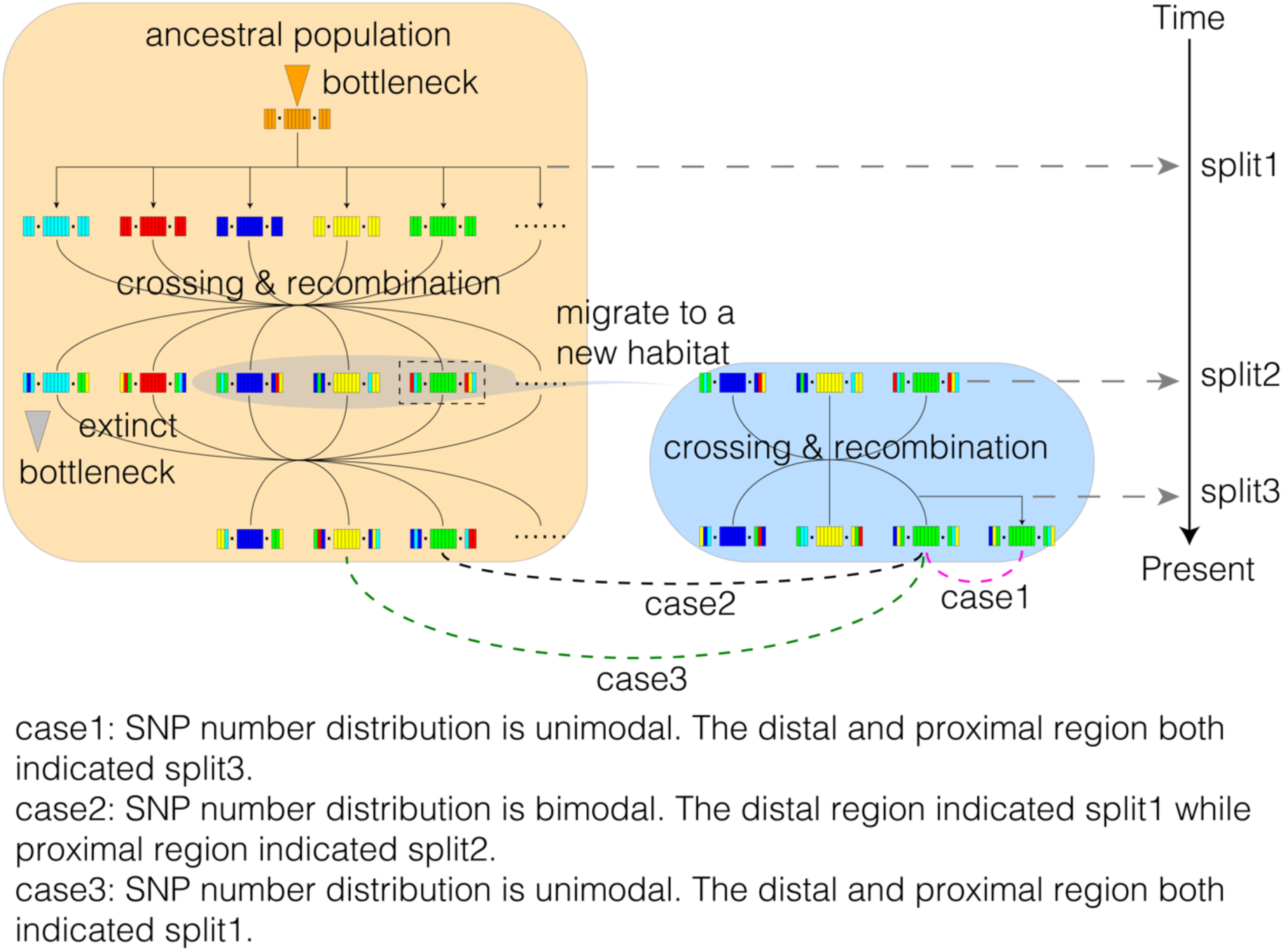
Patterns of sequence divergence differ along the genome. This figure complements **Extended Data Fig. 2b**. The three numbered cases shown in both figures correspond to each other. The groups of colored bars each represent a single chromosome of different wild barley individuals. The outer bars stand for genomic windows in distal regions, the inner ones for those in proximal regions. After an initial bottleneck an ancestral population of wild barley split into structured subpopulations (represented by different colors). This process may have occurred independently in different mutually isolated populations (e.g. in the Southern Levant or Central Asia). Here, one local ancestral population is shown as an example. Gene flow between subpopulations reshuffled haplotypes over time (left-hand part). As the population colonized new habitats, bottlenecks occurred ancestral haplotypes were lost (right-hand part). Since the recombination rate in proximal regions is low, these can be considered as a single recombinational unit (∼haplotype block). Because proximal regions are physically extensive, long shared haplotype blocks strongly influence the distribution of sequence divergence in windows of fixed physical size. In case1, two individuals are compared that come from the same extant wild barley population and neither has received recent gene flow from other populations. They share the same haplotype blocks in both distal and proximal regions and the distribution of sequence divergence is thus unimodal. In case2, two individuals come from a different population. Recombination and gene flow have reshuffled and broken up haplotypes in distal regions, whereas the proximal haplotypes trace back to a common ancestor at time point split2 and have remained intact in this scenario. Hence, the distribution of sequence divergence is bimodal: the peak in distal regions corresponds to early divergence (split1), whereas the peak in proximal regions reflects later divergence (split2) of the two lineages that lead to either individual. In case 3, also the proximal regions trace back to different ancestral haplotypes, which diverged early (split1). Hence, the distribution of sequence divergence along the genome is unimodal with a single early peak.

**Supplementary Figure 2:**
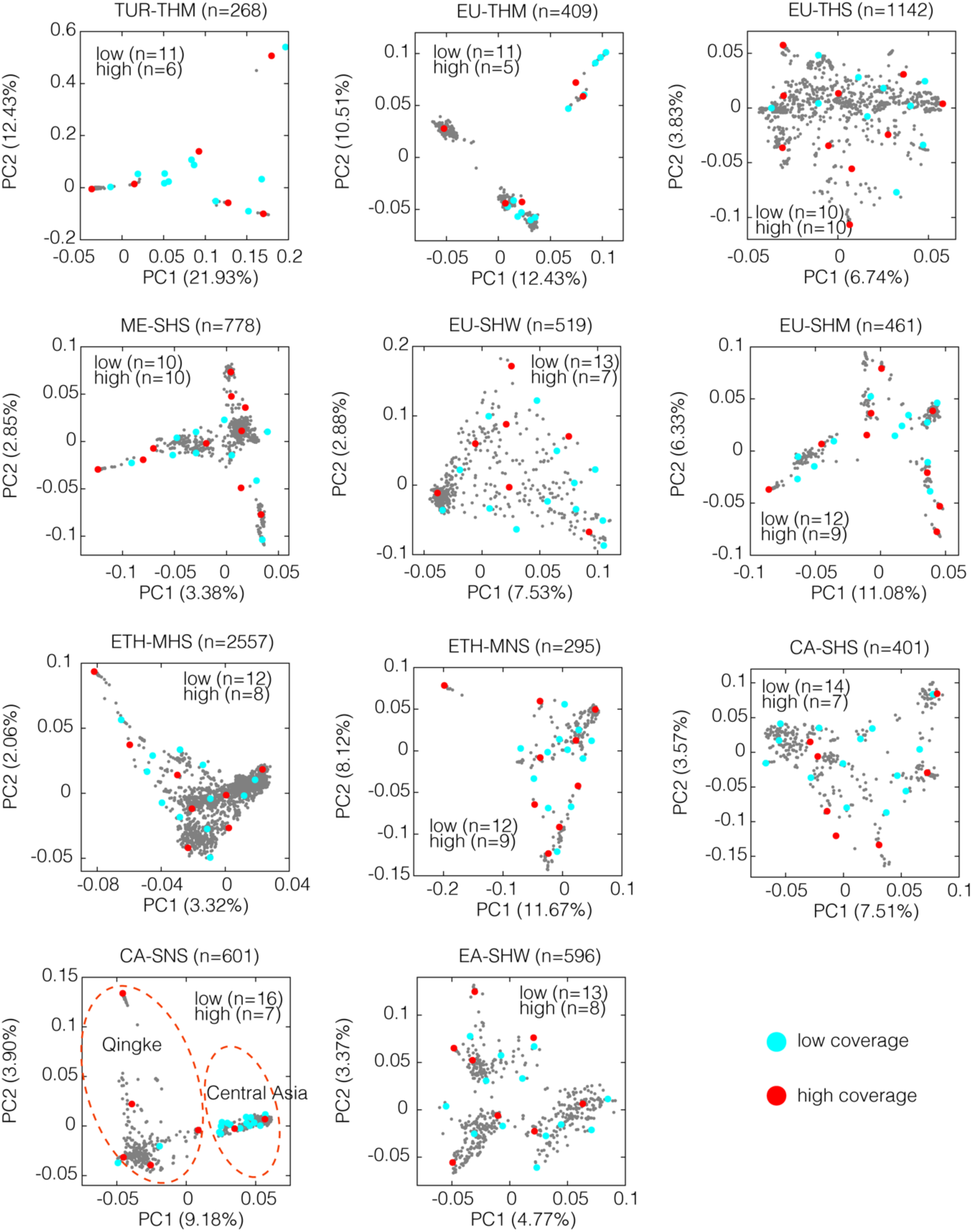
Selection of samples for whole-genome sequencing from 11 domesticated barley populations. Milner et al. 2019 defined 12 populations of domesticated barley. The selection of samples from their “orange” population is shown in **Supplementary Fig. 3**. For each of the remaining 11 populations, PCAs were run for un-admixed samples, i.e. those with ADMIXTURE ancestry coefficients ≥ 0.95. Then, about 20 samples (13 high-coverage, 7 low-coverage) were selected to cover the PCA diversity space of each population. The left-hand cluster in the PCA of CA-SNS (Central Asia 6-rowed naked barleys) contains Qingke (Tibetan hulless barleys). This population was studied by Zeng et al. 2018 in detail and was not the focus of the present study. Hence, comparatively fewer Qingke samples were selected.

**Supplementary Figure 3:**
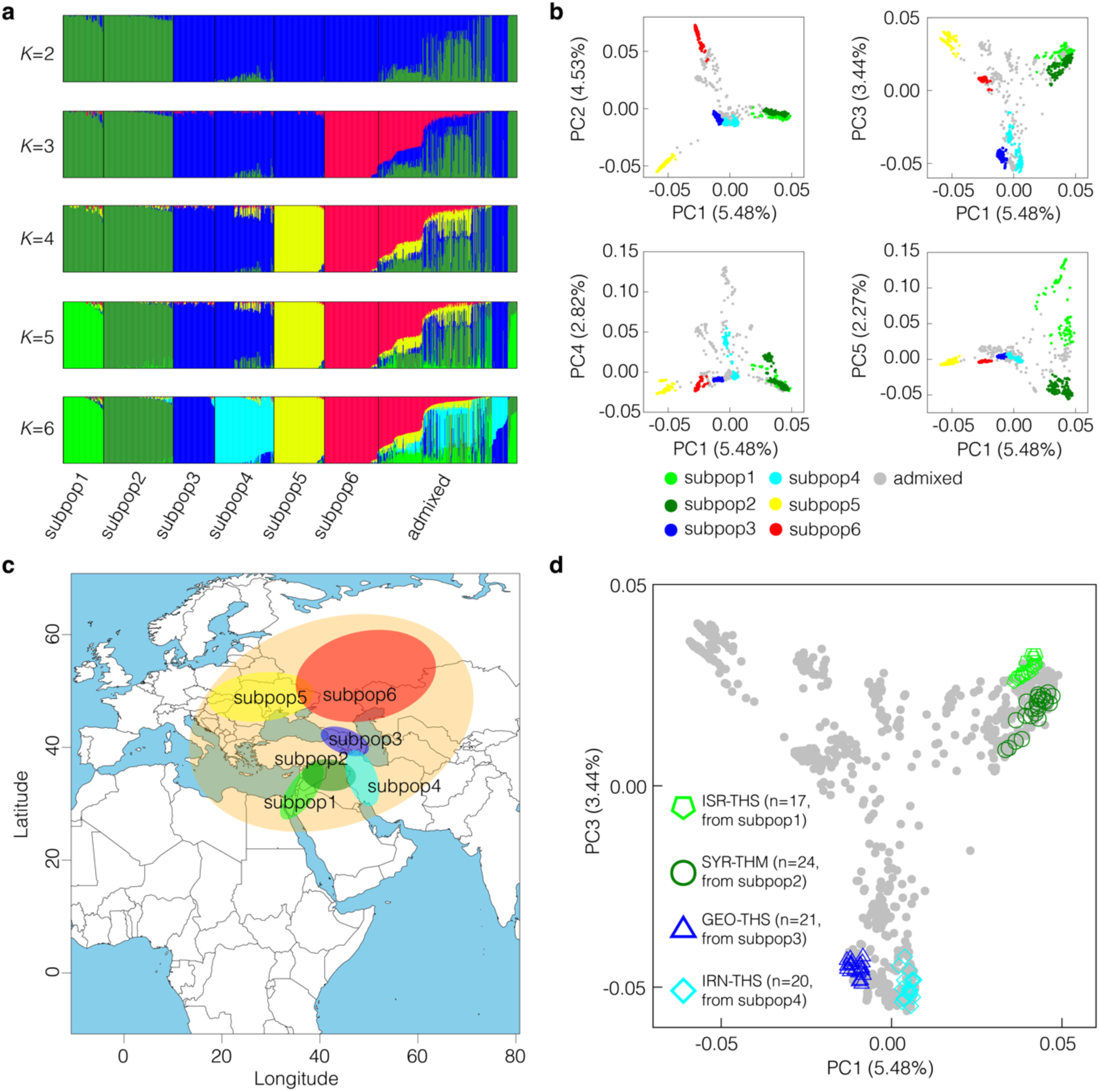
Selection of samples for whole-genome sequencing from the “orange” population of Milner et al. 2019. This figure is a complement to the sample selection in **Supplementary Fig. 2. (a)** Individual ADMIXTURE ancestry coefficients with the number of ancestral populations (*K*) ranging from 2 to 6. ADMIXTURE was run on member of the Milner et al.’s “orange” population using their GBS data. Individuals whose major ancestry coefficient was less than 0.8 were considered admixed. **(b)** Diversity space of the “orange” population as revealed by PCA. The higher PCs (2 to 5) were plotted against PC1. Colors correspond to unadmixed samples as per panel **(a)**. **(c)** Predominant geographical origins of the subpopulations of “orange”. **(d)** Samples from four populations in the Near East and Causasus that were selected for whole-genome sequencing are highlighted in the PCA.

**Supplementary Figure 4:**
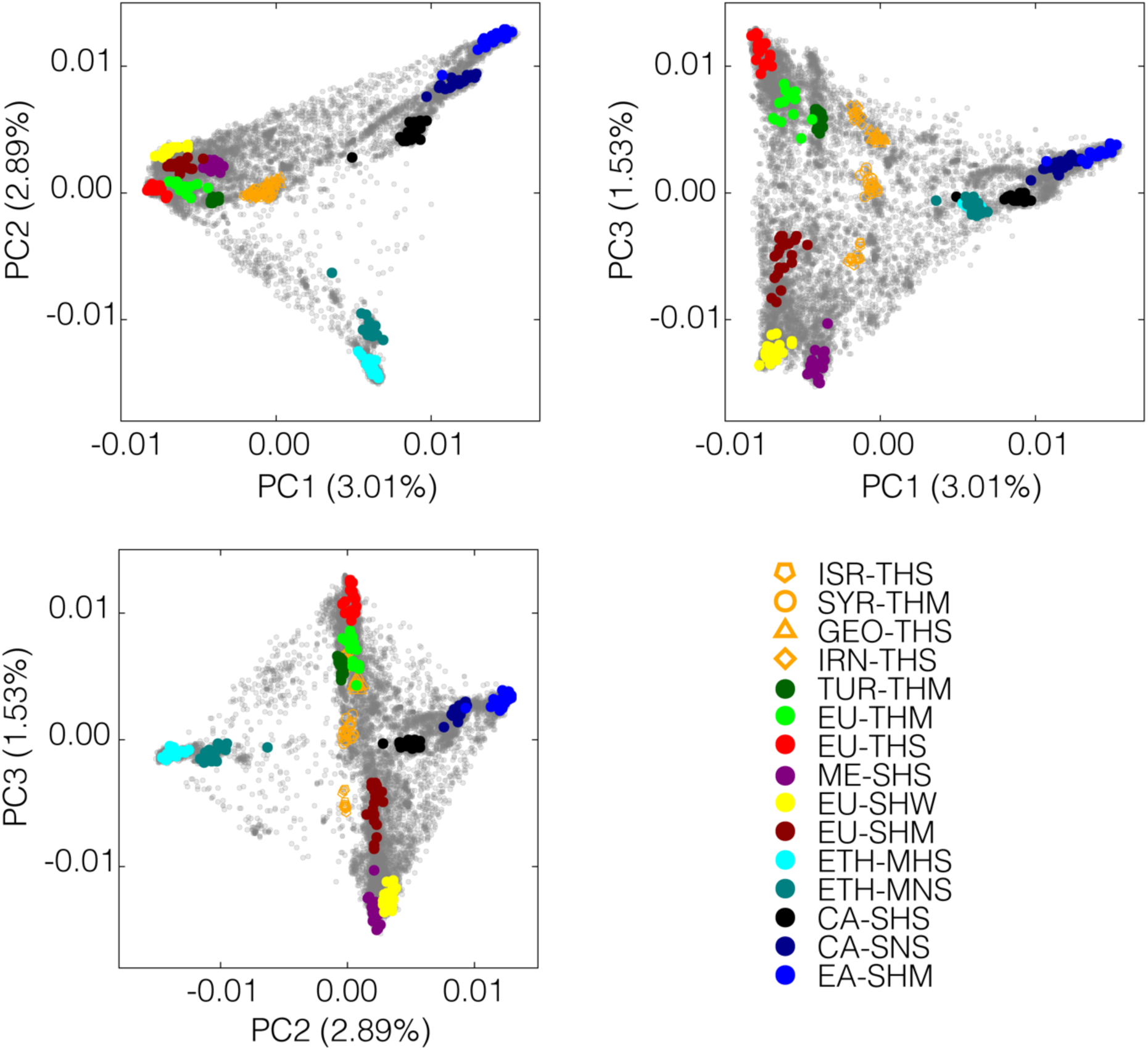
Positions of selected samples for whole-genome sequencing in the diversity space spanned by 19,778 domesticated samples of Milner et al. 2019.

**Supplementary Figure 5:**
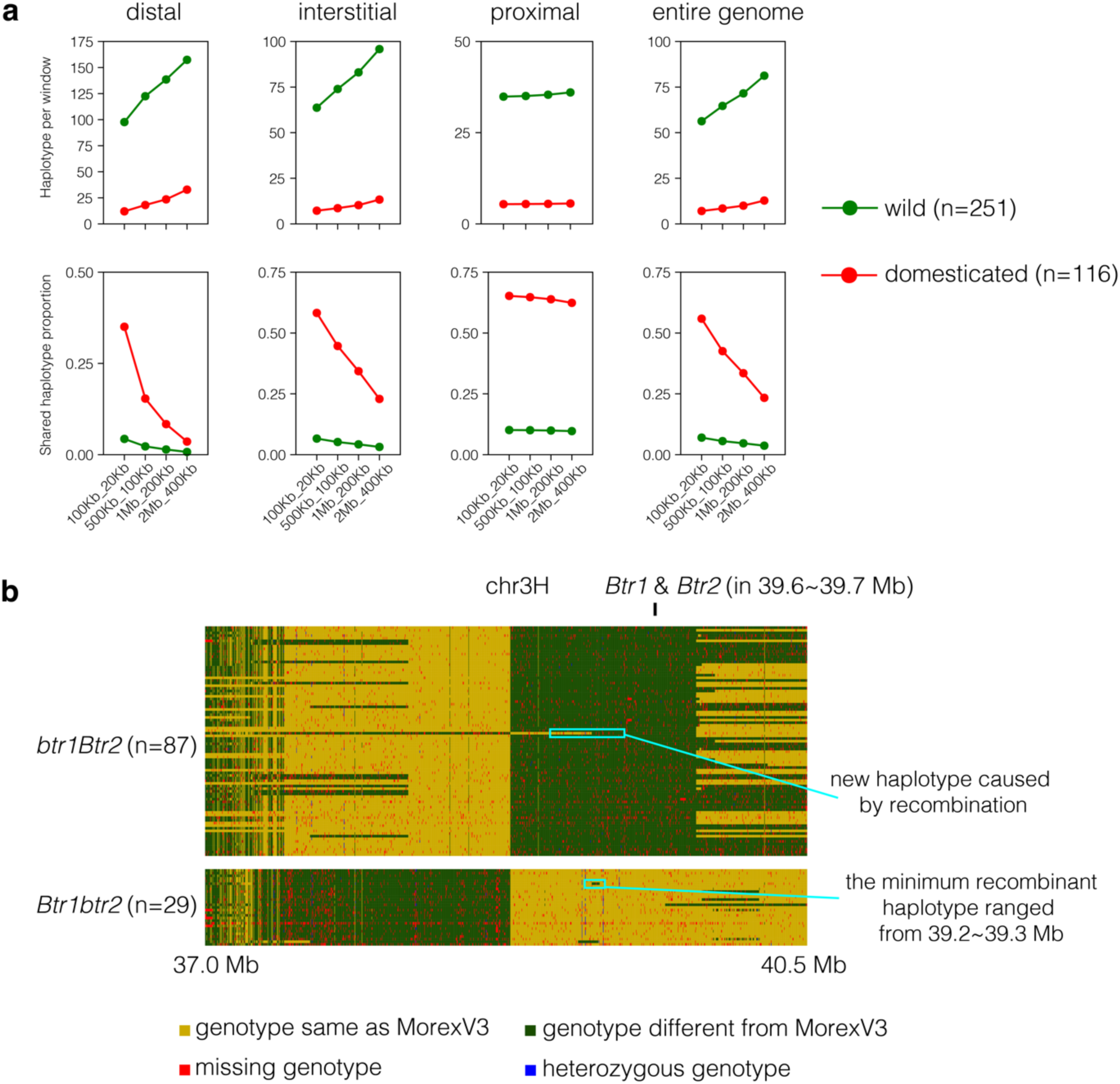
Effect of window size on haplotype definition. **(a)** Haplotype numbers and proportions of shared haplotypes in wild and domesticated barleys were computed from IntroBlocker runs with different window sizes: 100 kb (shift: 20 kb), 500 kb (shift: 100 kb), 1 Mb (shift: 200 kb), 2 Mb (shift: 400 kb). As the window size increases, so does the number of haplotypes, whereas the proportion of shared haplotypes decreases. The pattern is less pronounced in proximal, recombination-poor regions where haplotypes are longer. **(b)** The haplotype structure at the *btr1/2* locus is shown as an example. The samples shown are cultivated barleys with recombinant haplotypes. If the window size is large, more recombinant haplotypes are considered novel. The length of the smallest sequence exchange we observed between the original haplotypes (*Btr1btr2* and *btr1Btr2*) was 100 kb. Hence we used this window size (with a 20 kb shift) to compile the haplotype matrix for subsequent analyses.

**Supplementary Figure 6:**
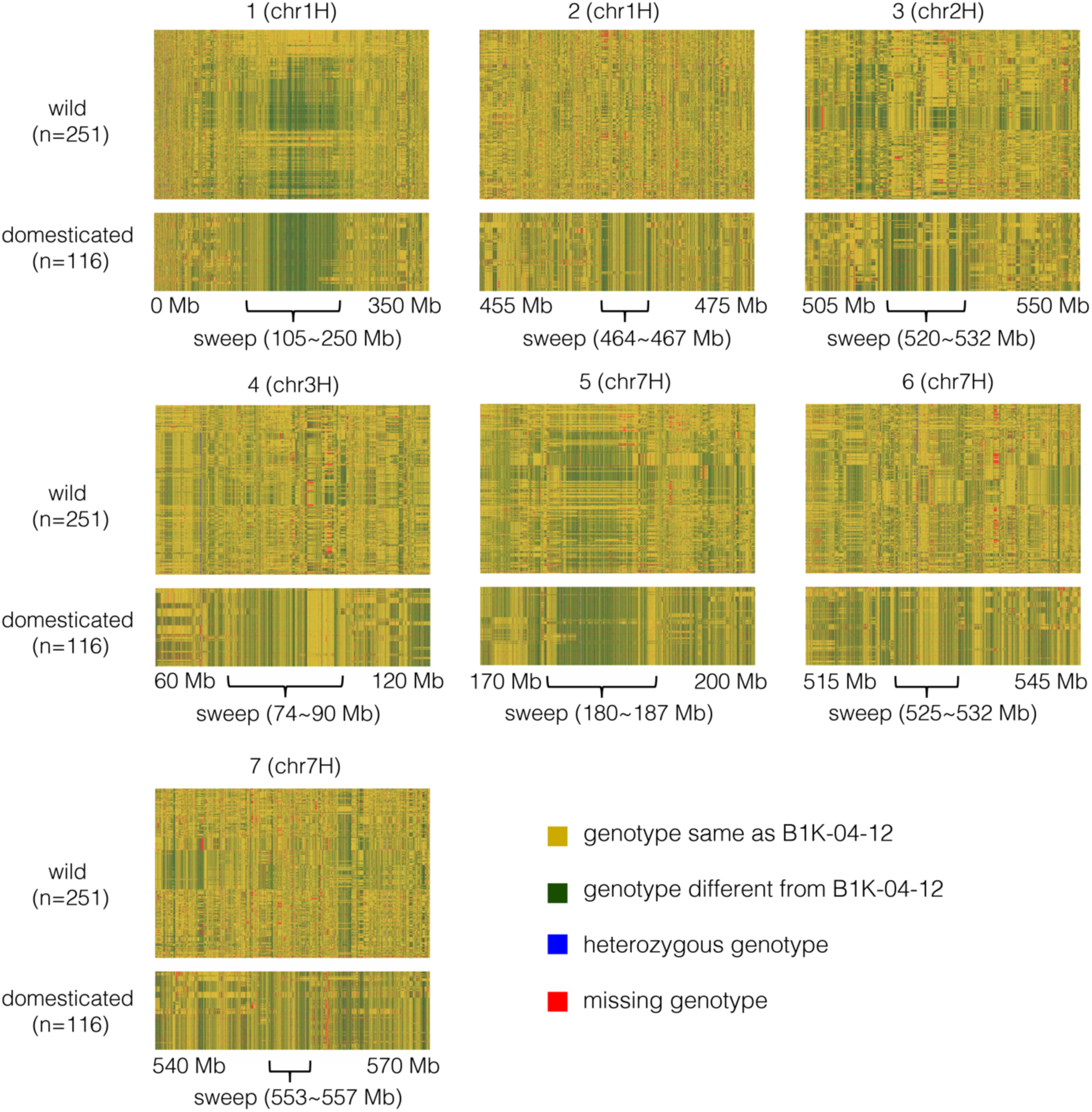
SNP haplotypes in the seven selective sweep regions shown in. Fig. 3g. The reference for SNP matrix was the wild barley B1K-04-12 from the barley pangenome (Jayakodi et al. 2023). Genotype calls were color-coded as indicated. Only SNPs sites with fewer than 20% missing calls and a minor allele frequency above 10 % were used.

**Supplementary Figure 7:**
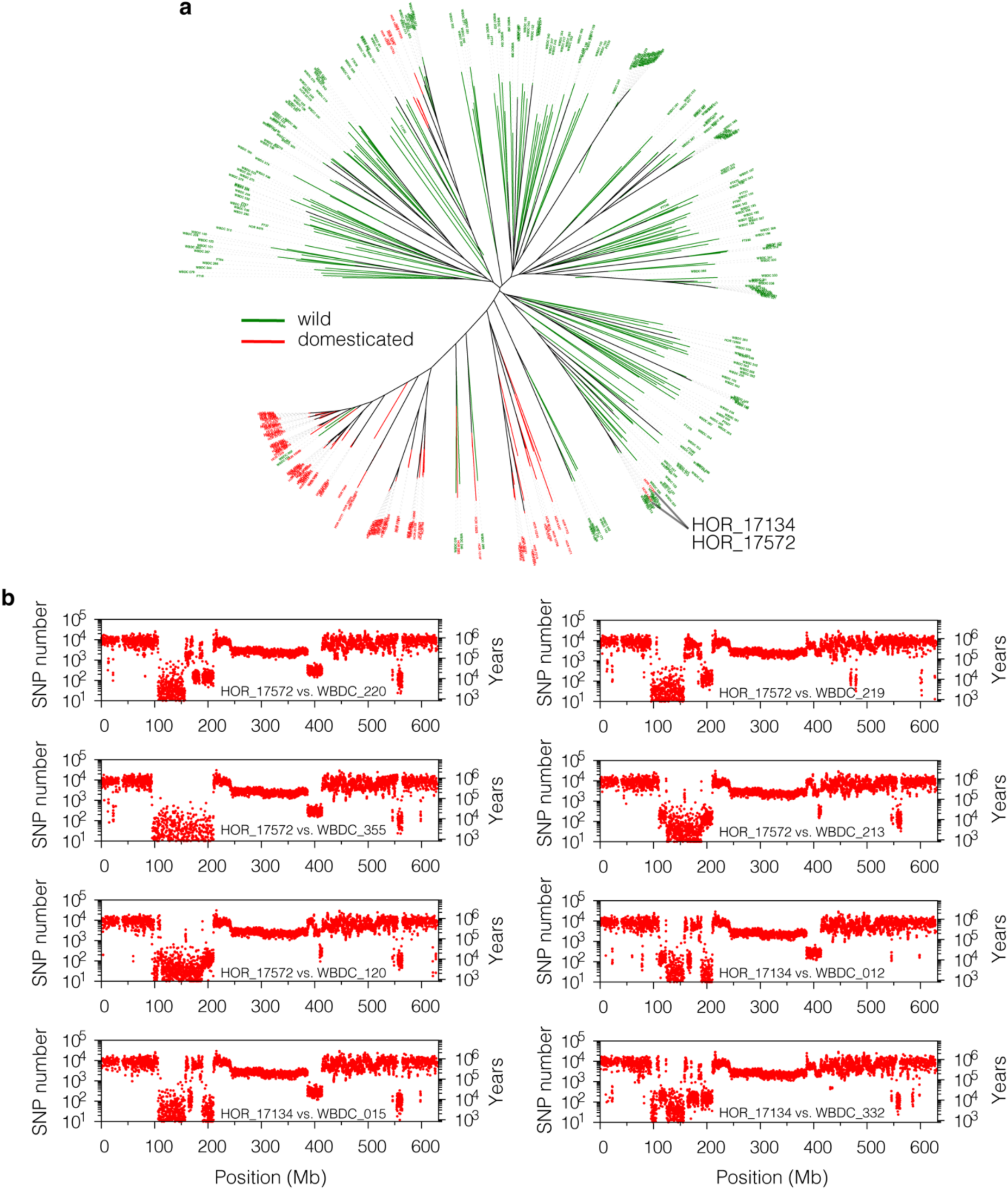
Recent gene flow from Eastern Asia wild barley to European 6-rowed winter barleys on chromosome 7H. **(a)** Neighbor-joining tree computed from 1.85 M biallelic SNPs in the interval 120 Mb to 160 Mb on chromosome 7H. Two accessions, HOR 17134 and HOR 17572, cluster with wild barleys. **(b)** Sequence divergence (SNPs per Mb) between these two accessions and different wild barleys in 100 kb windows (shift: 20 kb).

**Supplementary Figure 8:**
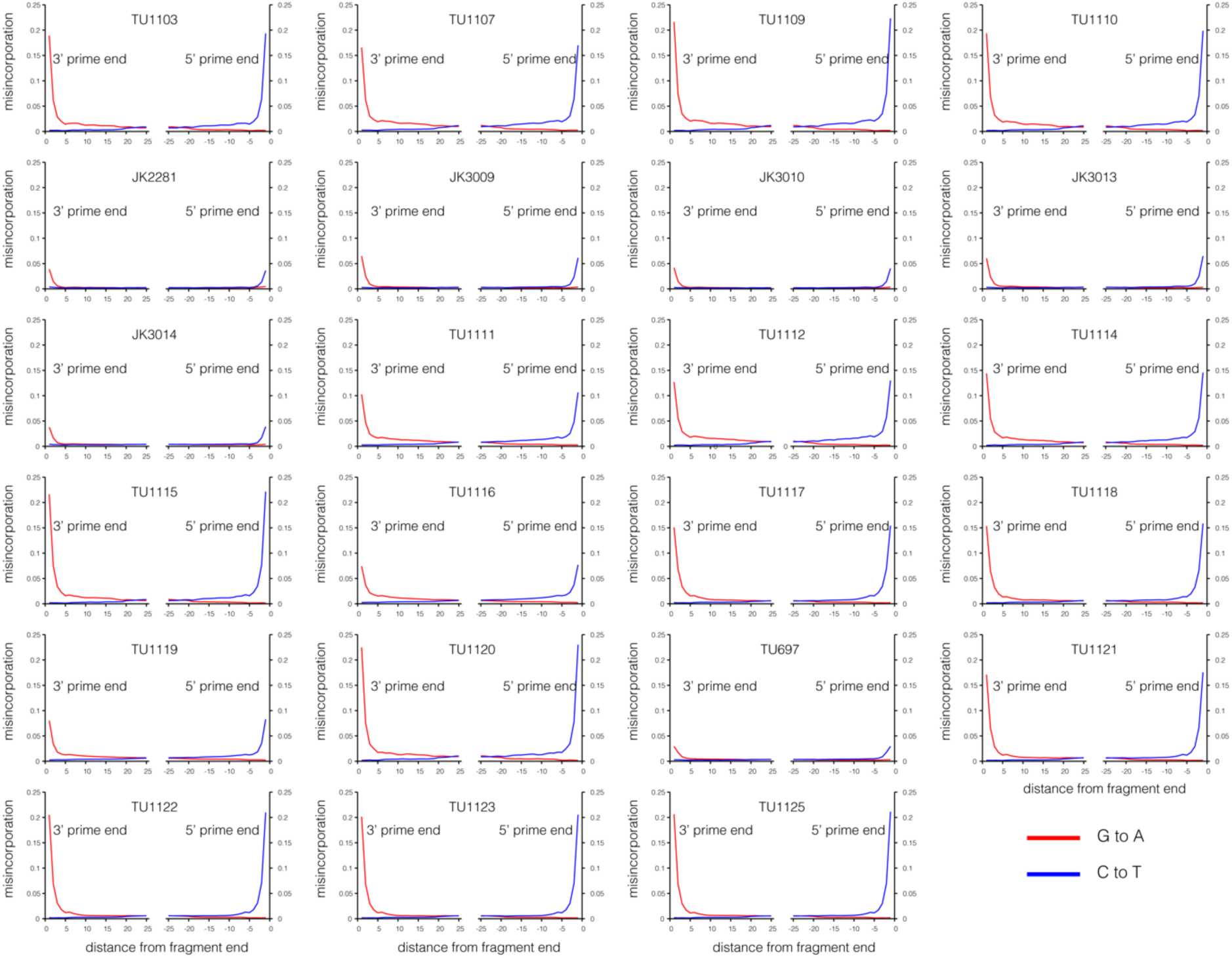
Nucleotide misincorporation profiles in the sequence data 23 ancient DNA samples.

**Supplementary Figure 9:**
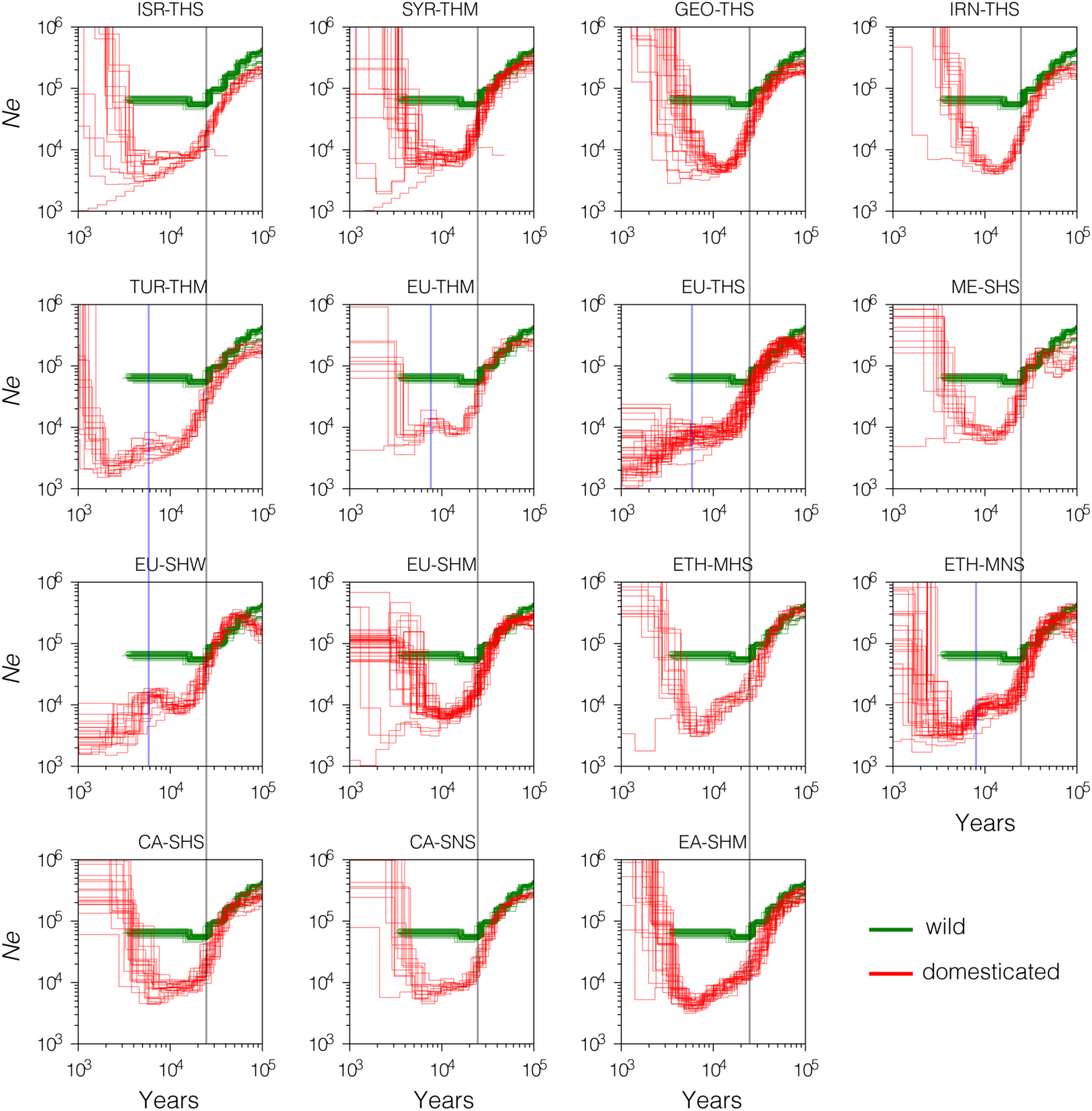
Historic trajectories of effective population sizes in wild and domesticated barley as inferred by PSMC. Each red line stand for one of pseudo-diploid genomes representing all pairwise combination of samples in the respective populations. The green lines (shared between panels) stand of the pseudo-diploid genomes derived from the sample pairs listed in **Supplementary Table 15.** The grey line marks the incipient divergence of wild and domesticated population starting about 25 ka BP. The blue line in some barley population marks a later bottleneck that coincides with the geographic range expansion of domesticated barley.

**Supplementary Figure 10:**
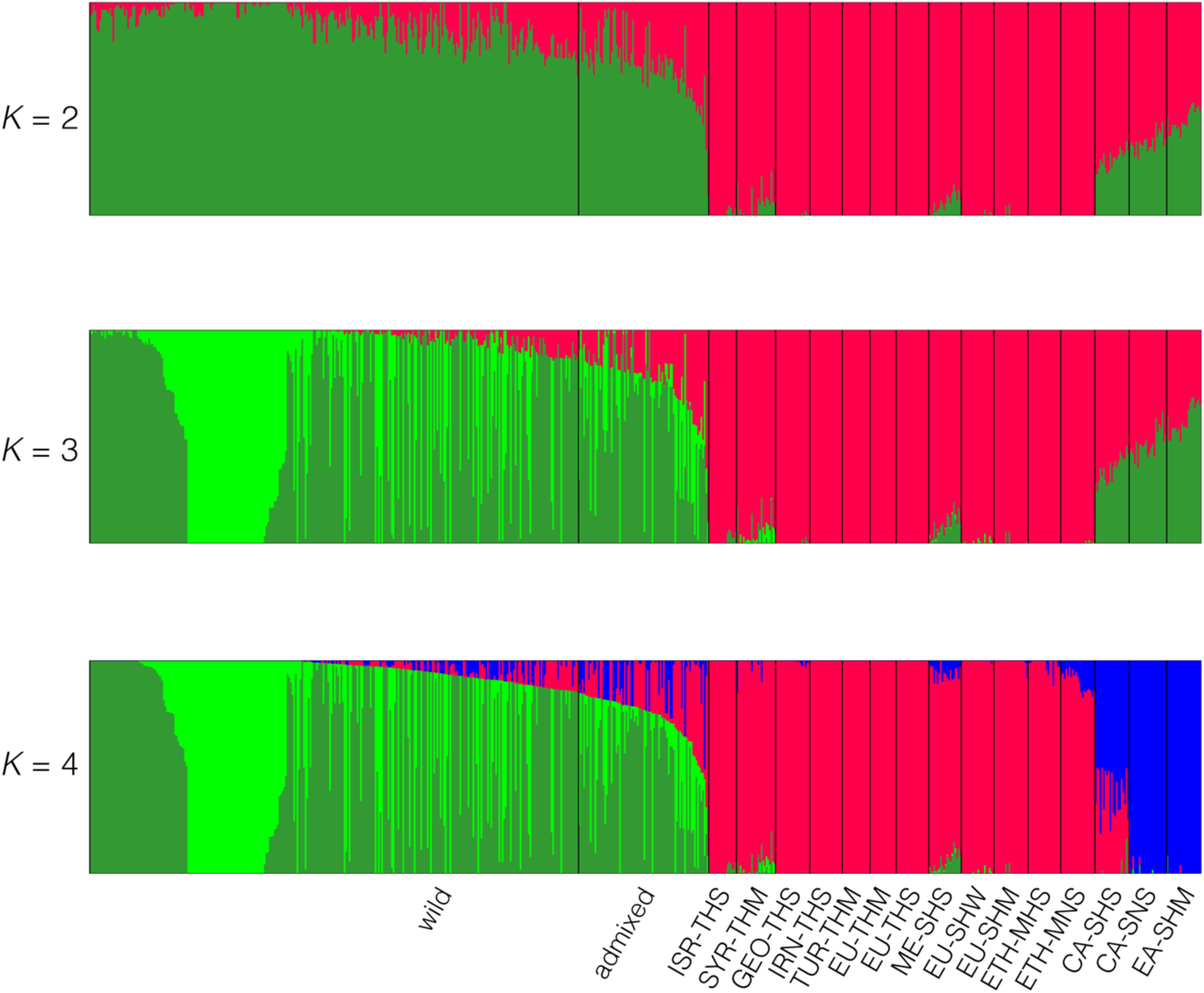
Individual ancestry coefficients in 682 barley samples as determined by ADMIXTURE with the number of ancestral populations (*K*) ranging from 2 to 4 for all 682 samples. Individuals designated as “wild” in the passport data but with <85% of “wild” ancestry (green color) were considered admixed and excluded from subsequent analyses.

**Supplementary Figure 11:**
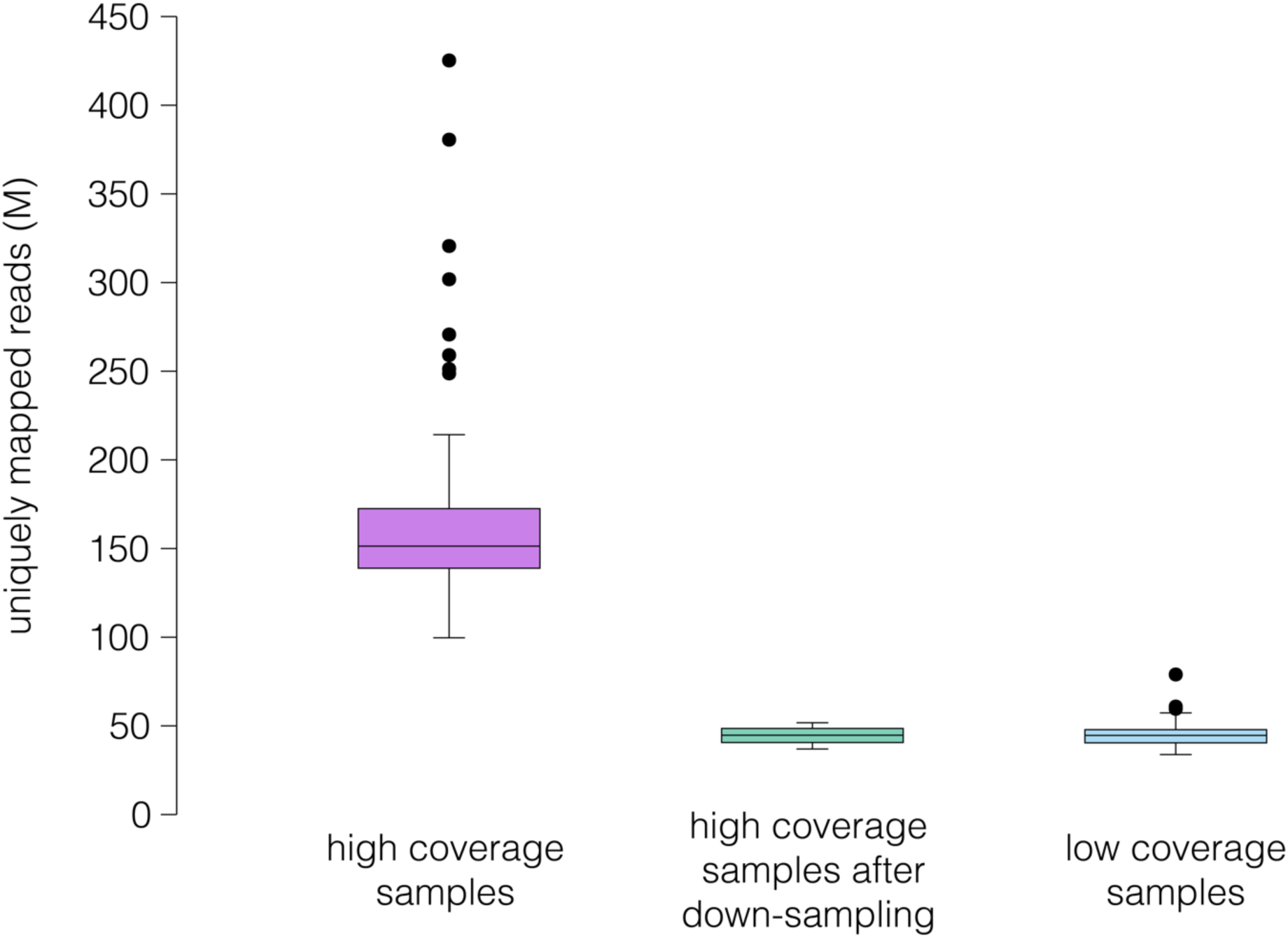
Down-sampling of high-coverage sequence data. Box plots showing the distribution of the numbers of uniquely mapped (MAPQ ≥ 20) reads for 116 high-coverage and 186 low-coverage samples. For some analyses, high-coverage data were down-sampled to the level of the low-coverage samples.

**Supplementary Figure 12:**
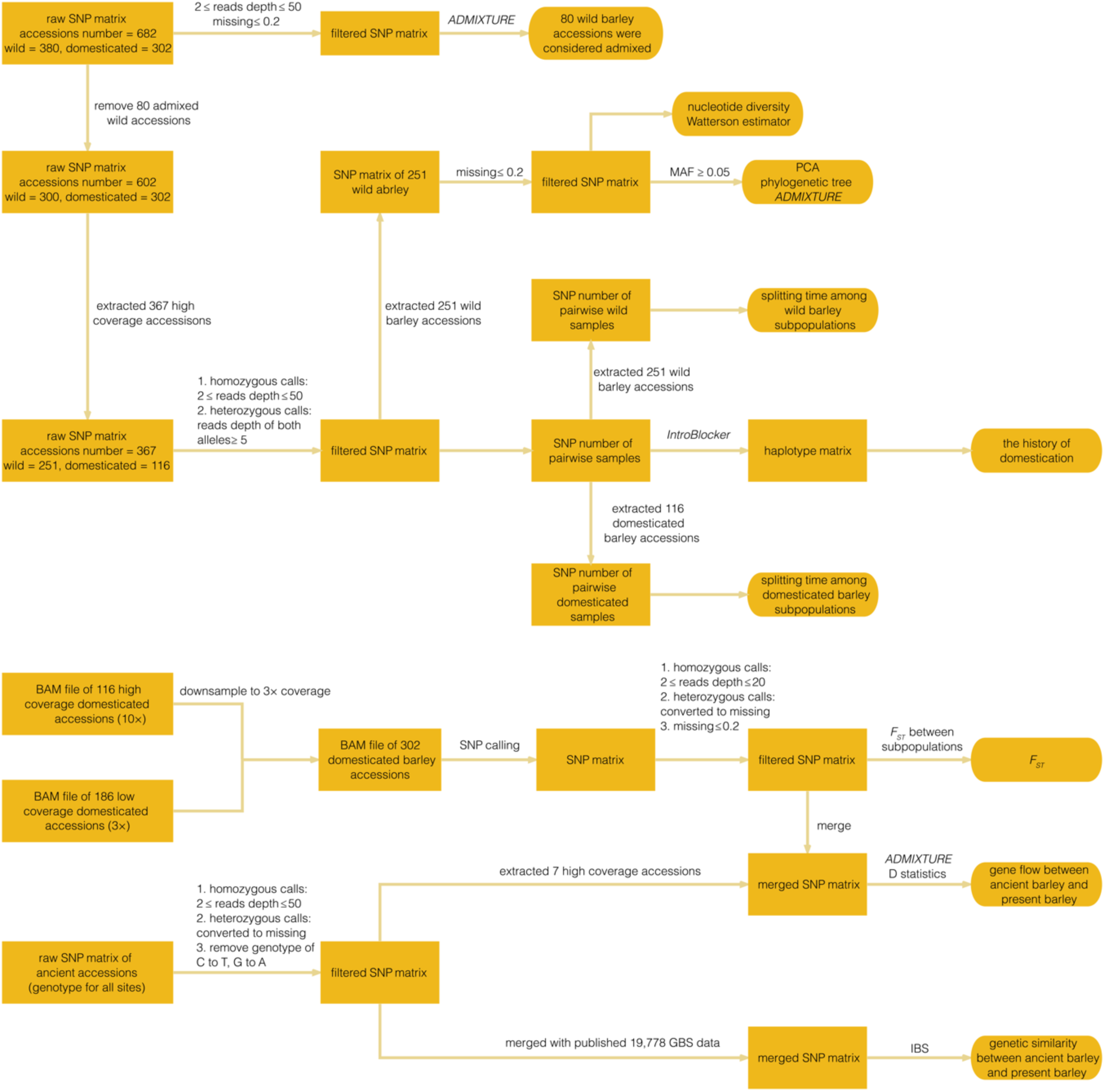
Workflow of analyses conducted on SNP matrices.

**Supplementary Figure 13:**
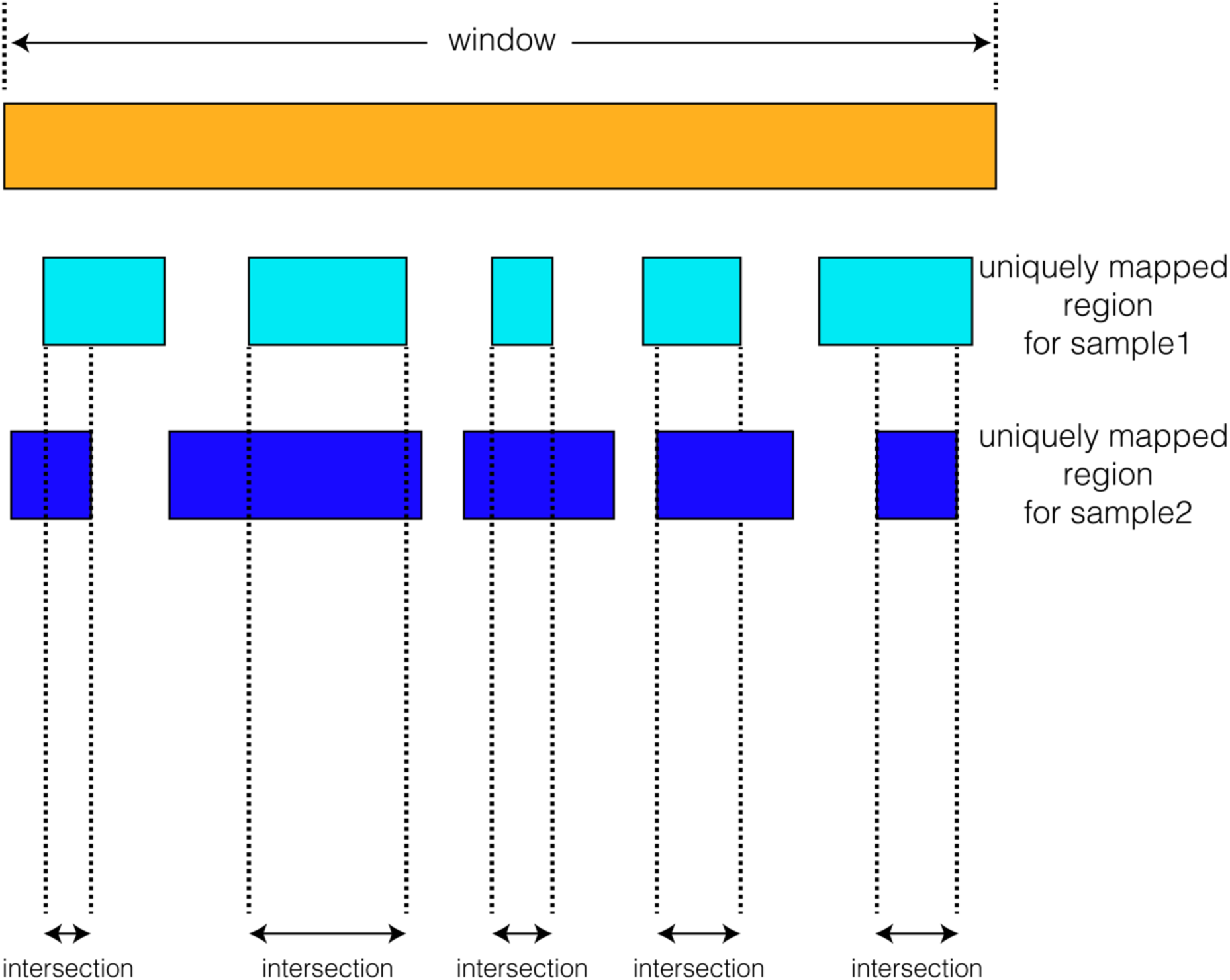
Normalization of SNP counts for read depth. To determine the number SNPs between two samples in a given genomic window of size, we first intersect the regions covered by at least two uniquely mapped (MAPQ ≥ 20) reads. Then, we calculated the normalized SNP number according to the formula: (raw SNP number / cumulative of the intersected intervals) × window size. The “raw” SNP number was determined with command “sample-diff counts-only” of Plink2.

**Supplementary Figure 14:**
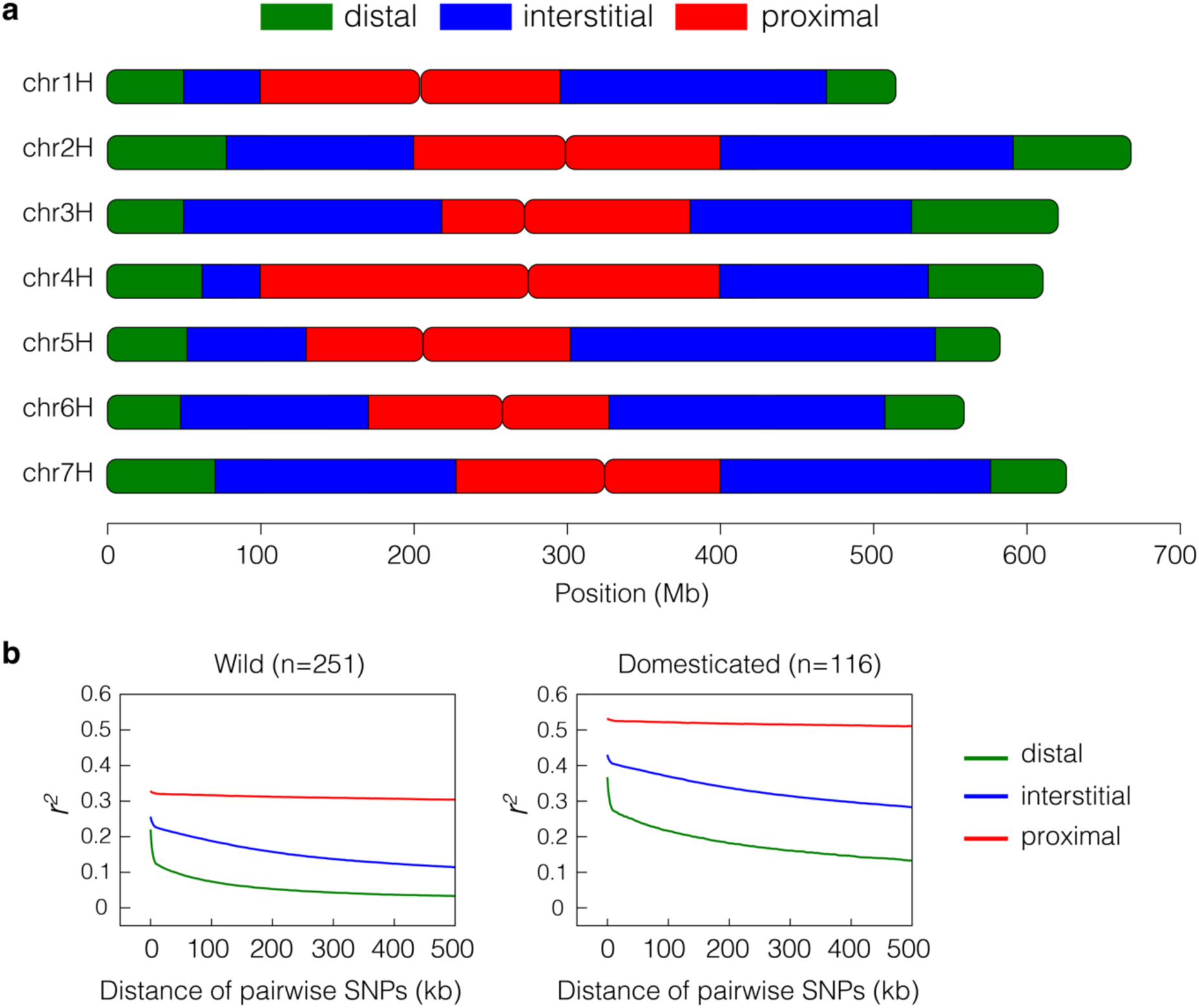
Decay of linkage disequilibrium (LD) in three genomic compartments. **(a)** The distal, interstitial and proximal regions were defined based on differences in recombination rate following Mascher et al. 2017. The precise boundaries of the three compartment of each chromosome are listed in **Supplementary Table 14**. **(b)** LD decay in the three genomic compartments thus defined in wild and domesticated barley.

**Supplementary Figure 15:**
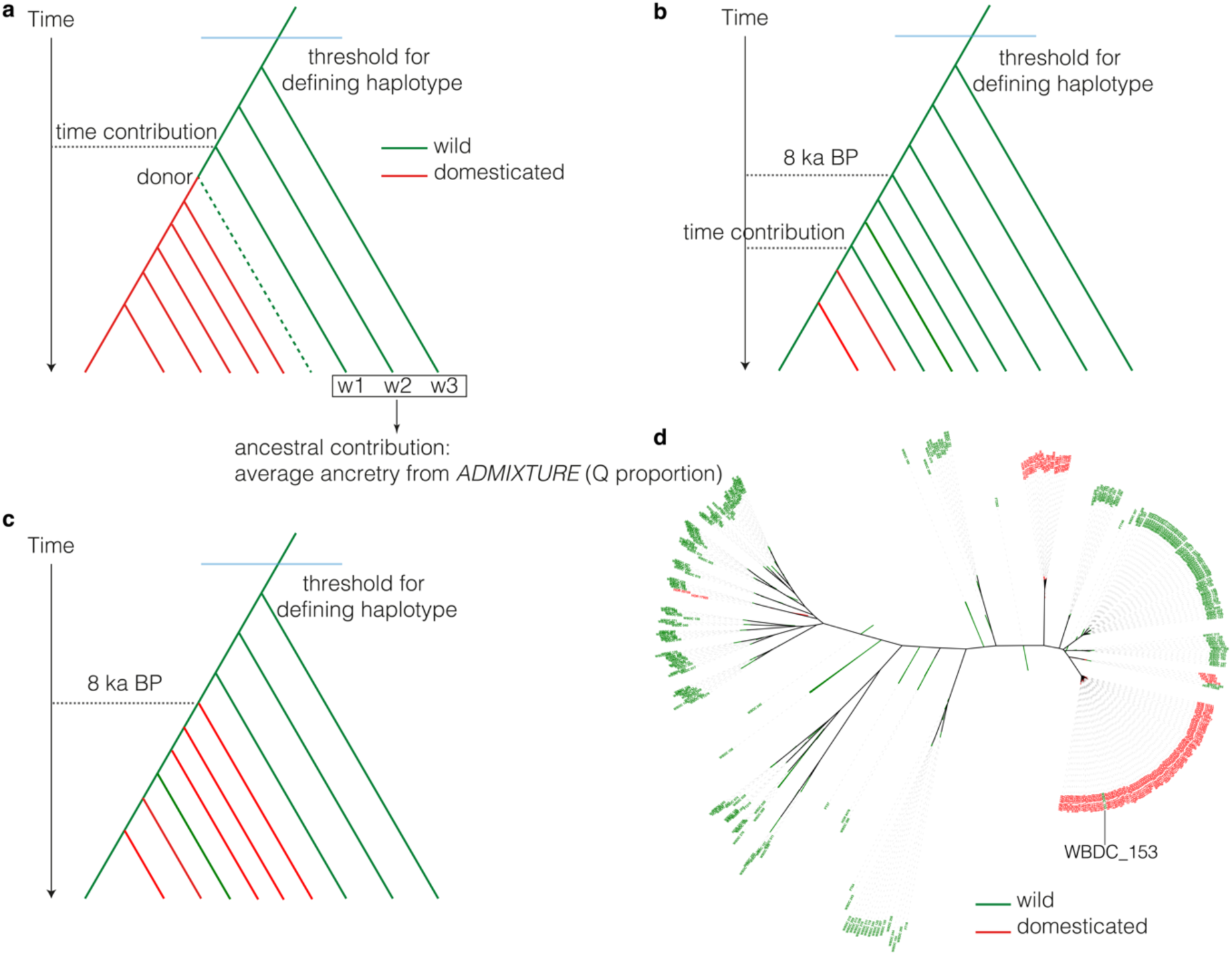
Inference of haplotype origins in time and space. Samples in trees have the same haplotype (H) in a genomic region of interest, i.e. their sequence divergence is below a chosen threshold. Red and green color stand for domesticated and wild samples, respectively. **(a)** Dating in the absence of post-domestication introgression. The time of origin of H was set to set time of divergence of the domesticated branch from the closest wild sample (w1). To find the most closely related wild sample, we compared the results of IntroBlocker runs with different thresholds: 400 SNPs (equivalent to an approximate divergence time of 32,000 years ago), 98 SNPs (8,000 years), 73 SNPs (6,000 years), 49 SNPs (4,000 years), and 24 SNPs (2,000 years). To determine the wild source population of population, the ancestry coefficients of all wild barleys (w1, w2, w3) with haplotype H were averaged. In panels **(b)** and **(c)**, the divergence time between domesticated and wild carriers of H is less than 8,000 years, indicating recent gene flow, either in the direction wild > crop **(b)** and crop > wild **(c)**. **(d)**. Neighbor-joining tree constructed from SNPs on chromosome 4H, 250 Mb – 300 Mb. The sample WBDC 153 is the only wild barley that shared a haplotype with domesticated samples. The most likely explanation is geneflow in the direction crop > wild.

**Supplementary Figure 16:**
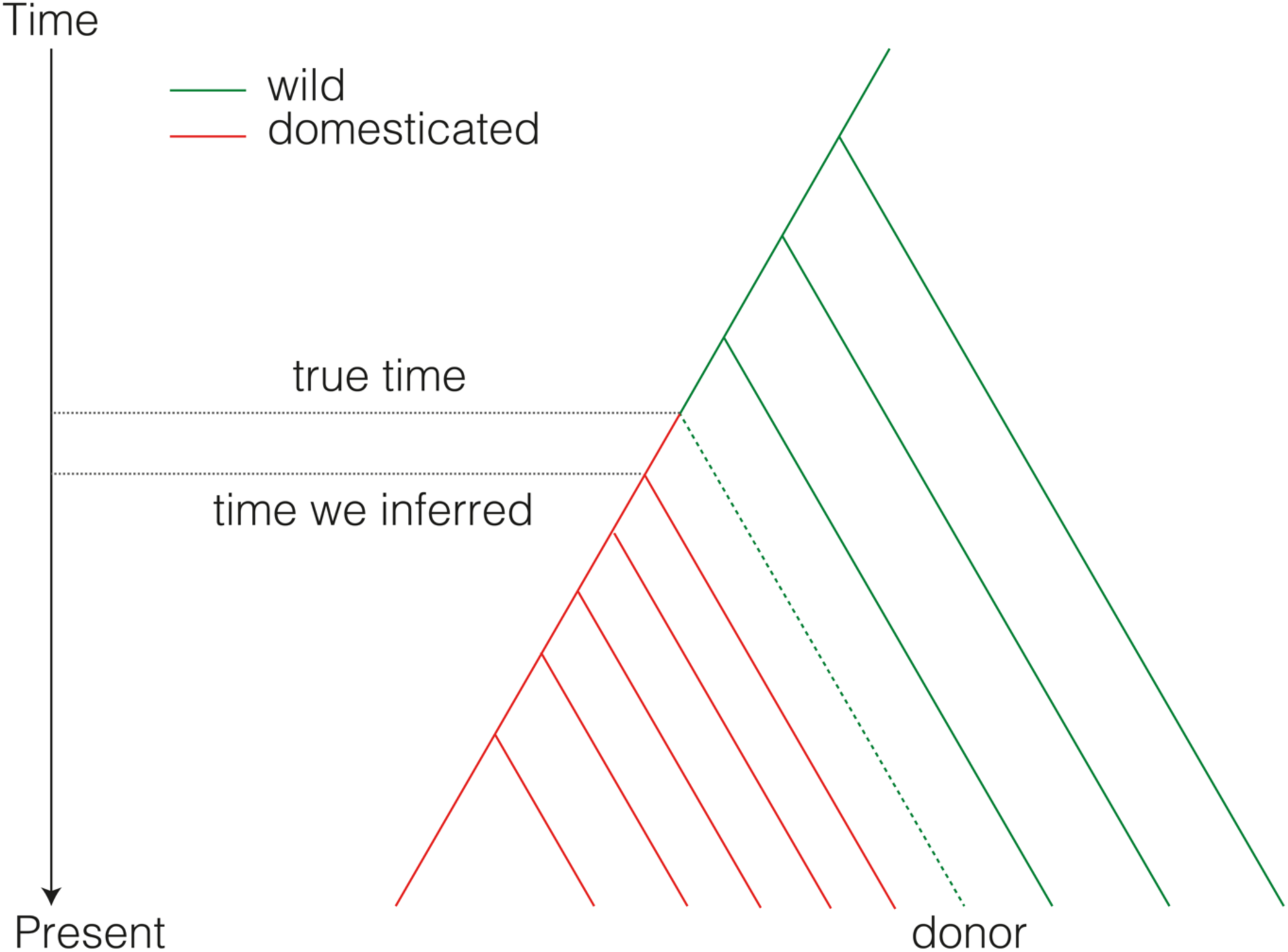
Dating the origin of haplotypes around domestication genes. This schema explains how we dated the emergence of mutations in single genes in **Extended Data Figure 9a**, which shows the distributions of sequence divergence (SNPs per Mb) in pairwise comparisons between all domesticated barleys at a locus of interest. The time inferred marks the earliest divergence between extant domesticated samples, but true time must be predate our estimate because the wild lineage (donor) in which the mutation arose is not represented in our data, for example because it became extinct or so rare as to avoid sampling.

## Extended Data Figures

**Extended Data Figure 1:**
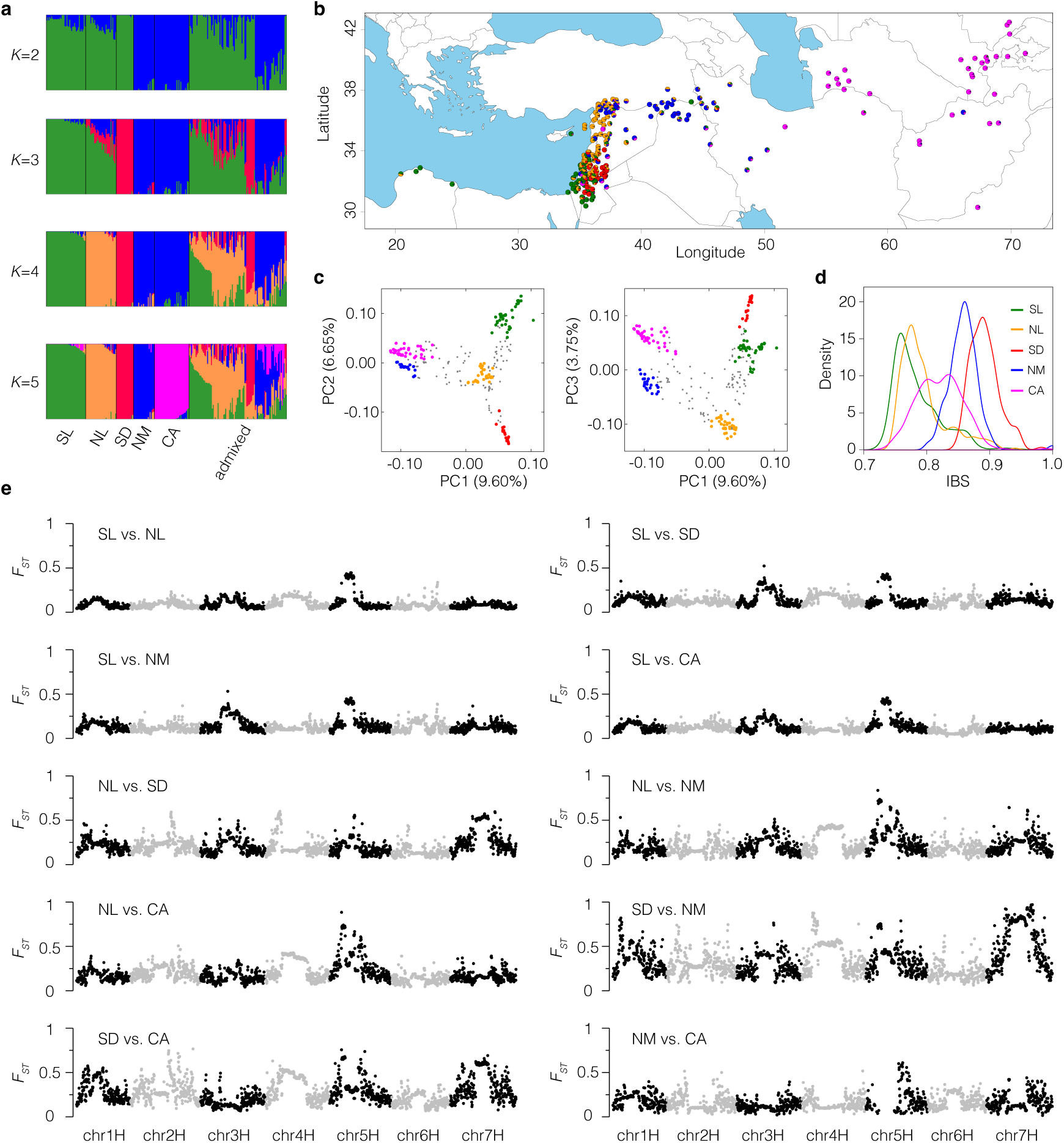
Population structure of wild barley. **(a)** Individual ancestry coefficients in ADMIXTURE with the number of ancestral populations (K) ranging from 2 to 5. Individuals with ≥ 85% ancestry form were considered un-admixed samples. **(b)** Collection sites of 237 wild barley samples with precise geographic locations. Pie charts show ADMIXTURE ancestry coefficients. **(c)** Principal component analysis (PCA) based on 37.14 million biallelic SNPs with a MAF > 5%. The first three PCs are shown (PC1 vs. PC2 and PC1 vs. PC3). Unadmixed samples are colored to their major ancestry component in panel **(a)**. **(d)** Distribution of pairwise identity-by-state (IBS) in the five wild barley populations. **(e)** Fixation indices (F_ST_) in 1 Mb windows along the genome. All possible contrasts between any two of the five wild barley populations are shown.

**Extended Data Figure 2:**
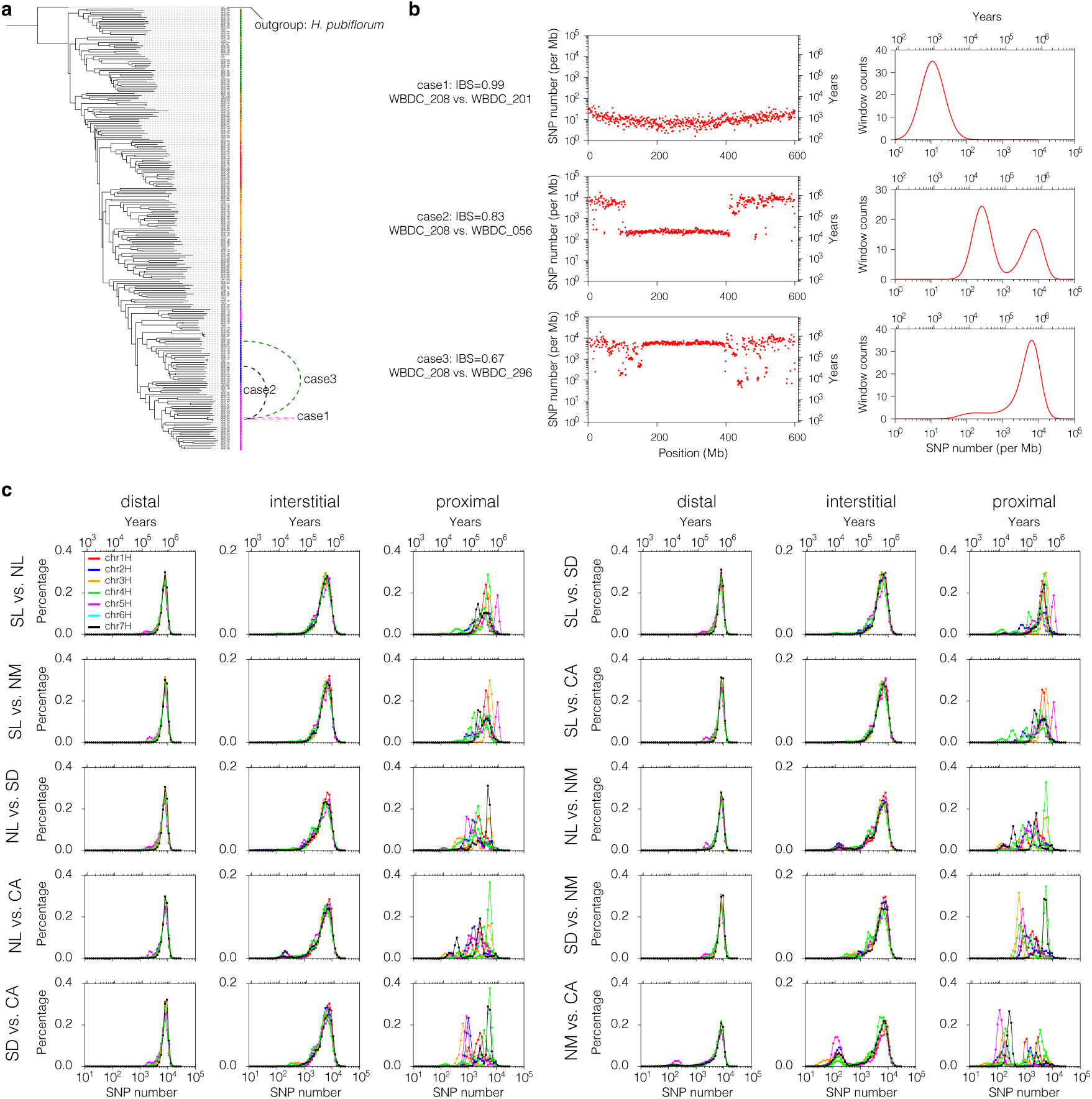
Sequence divergence between wild barleys in different genomic compartments. **(a)** Neighbor-joining tree of 251 wild barley genotypes computed from 37.14 M biallelic SNP markers with a MAF > 5%. *H. pubiflorum* was used an outgroup. Pie charts to the right show ancestry coefficients as determined by ADMIXTURE **(Extended Data Fig. 1)**. **(b)** The left-hand panels show the sequence divergence (SNPs per non-overlapping 1 Mb genomic windows) on chromosome 4H between three pairs of wild barleys. Case 1 each compares two members from the same population (CA, WBDC208 vs. WBDC201). In cases 2 and 3, WBDC 208 is compared to two members of the NM population, WBDC296 and WBDC056. The right-hand panels show the respective distribution densities. **(c)** Distributions of sequence divergence (SNPs per Mb) in three genomic compartments in pairwise comparisons between members of different wild barley populations.

**Extended Data Figure 3:**
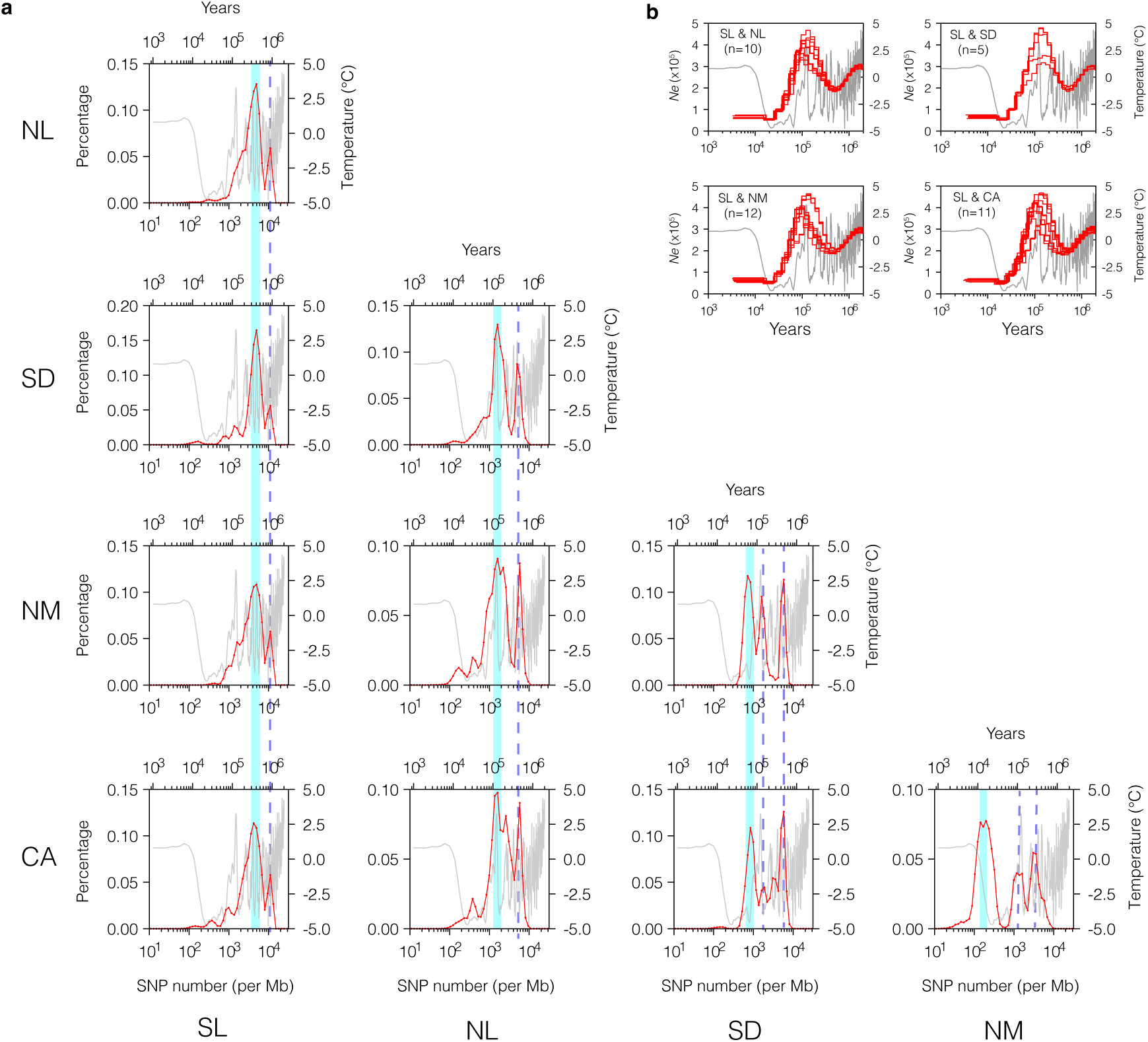
Demographic history of wild barley populations. **(a)** Distributions of pairwise sequence divergence (SNPs Mb) in pericentromeric regions (centromere +/- 25 Mb). The distributions were computed from comparisons between all possible combinations between members of the respective populations. Cyan shading marks the most recent peaks in the distributions interpreted as the most recent divergence from common ancestors. Dashed lines mark earlier such events. **(b)** Historic trajectories of effective population sizes computed with PSMC from pseudo-diploid genomes uniting members from different wild barley populations. One haploid genome was taken from the SL population, the other one from one of the four other populations. The number of pseudo-diploid genomes (n) for each population pair is indicated. Gray lines in panels **(a)** and **(b)** show the global average surface temperatures^39^.

**Extended Data Figure 4:**
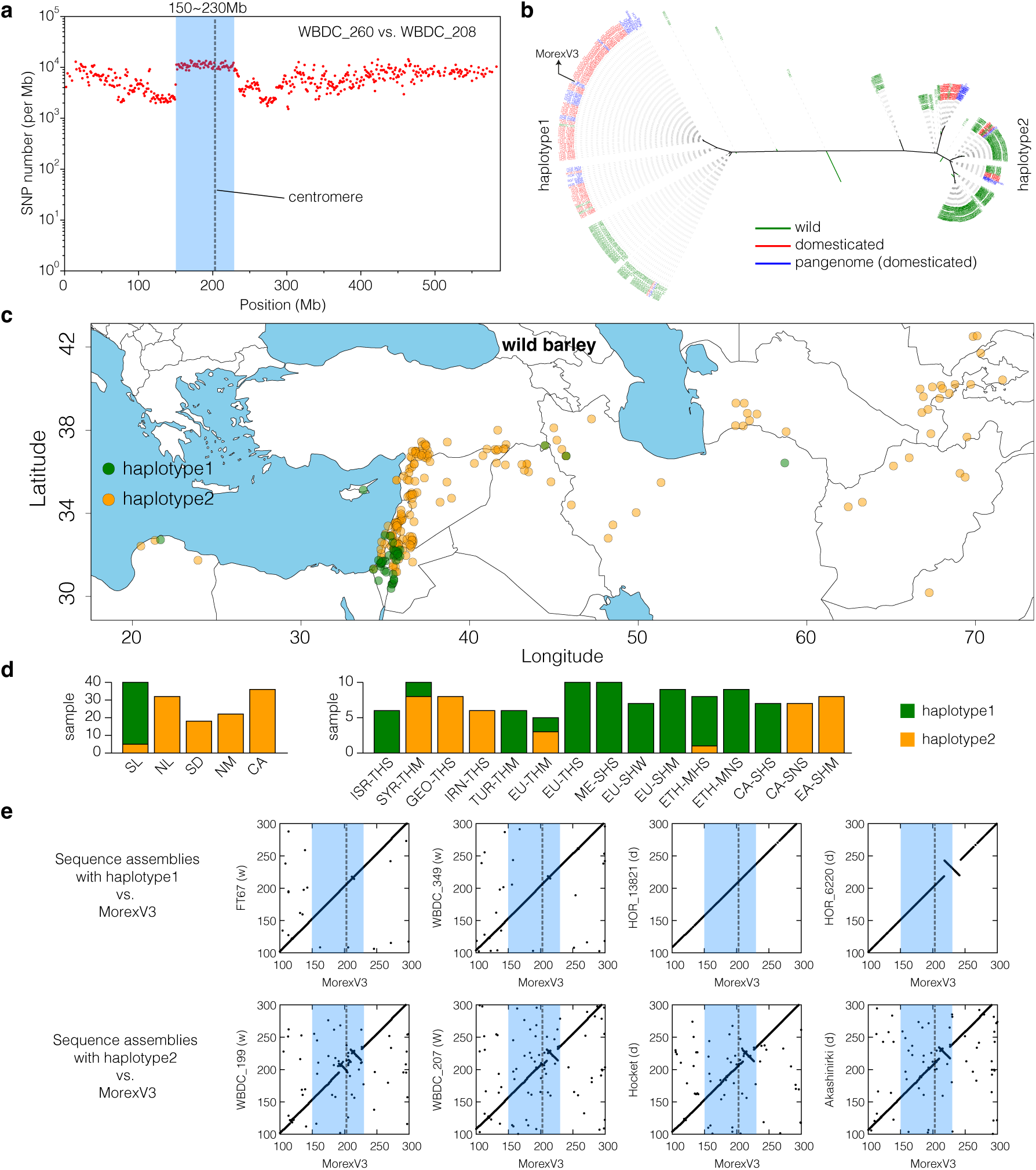
Deeply diverged pericentromic haplotypes on chromosome 5H. **(a)** Sequence divergence (SNPs per Mb) on chromosome 5H between wild barleys WBDC260 (SL) and WBDC208 (CA). **(b)** Neighbor-joining tree of wild and domesticated barleys including 48 samples from the pangenome of Jayakodi et al.^41^ **(c)** Collection sites of wild barleys carrying either haplotype. The “green” haplotype is most common in the SL population. **(d)** Frequencies of both haplotypes in wild and domesticated barley populations. **(e)** Alignments on chromosome 5H, 100 to 300 Mb between sequence assemblies of wild (w) and domesticated (d) pangenome accession with either haplotype to the MorexV3 reference. The accession names are indicated on the y-axis. In panels **(a)** and **(e)**, the boundaries of divergent haplotypes (150 to 230 Mb) are marked by blue shading and the dashed line at 205 Mb indicates the position of the centromere in the MorexV3 reference.

**Extended Data Figure 5:**
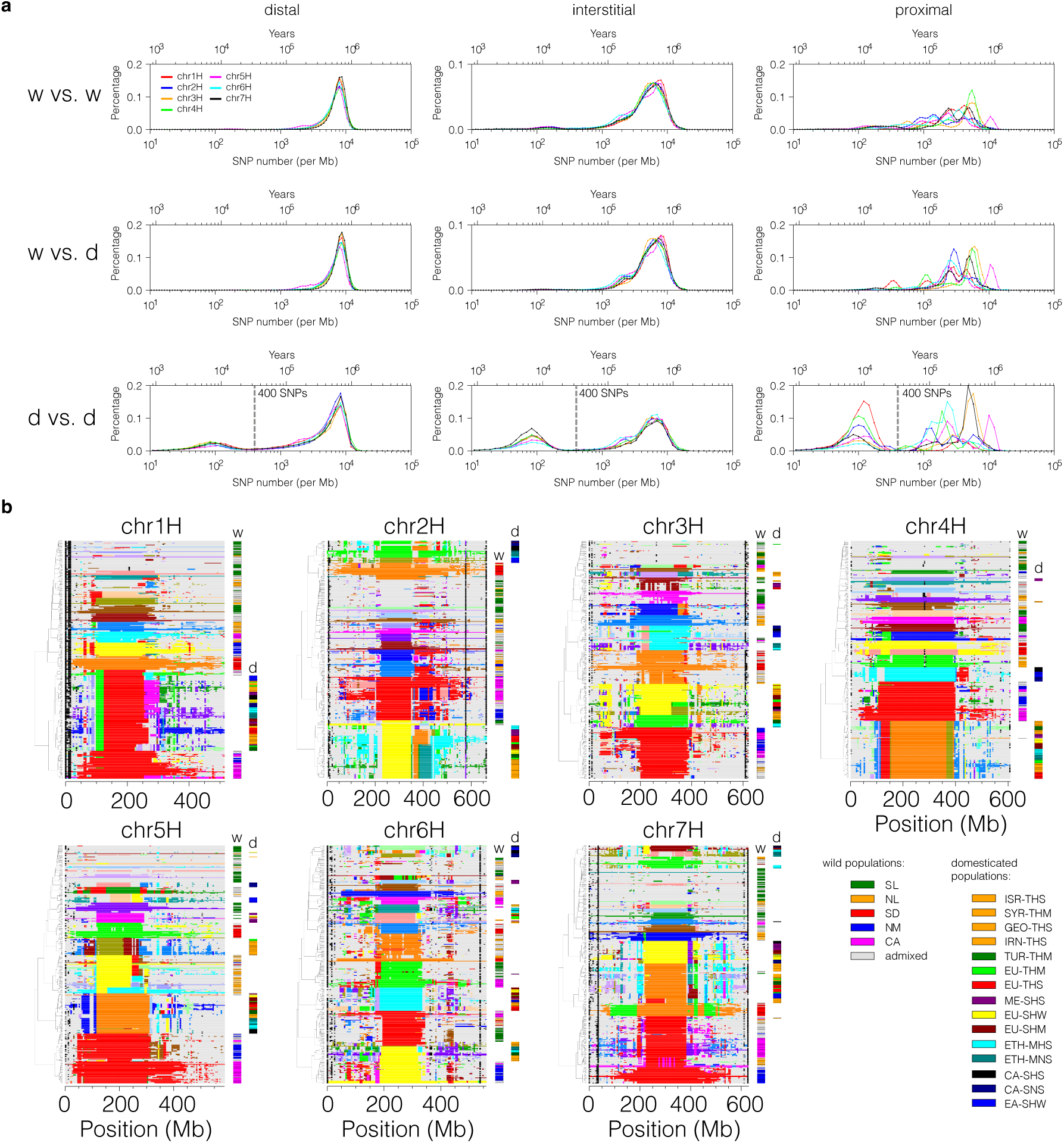
Ancestral haplotype groups (AHGs) in domesticated barley. **(a)** Finding a threshold for defining AHG. Sequence divergence (SNPs per Mb) in pairwise comparisons between wild (w) and (d) domesticated barleys. A value of 400 SNPs per Mb was chosen as the threshold to separate haplotypes in IntroBlocker. **(b)** Mosaic view of AHGs on the seven chromosomes of barley. The data shown are from an IntroBlocker run with a 5 Mb window size (shift: 5 Mb). Colors were assigned to 20 most frequent AHGs by IntroBlocker in semi-supervised mode giving priority to wild over domesticated samples. Black color indicates missing data, gray stands for less frequent haplotypes. Colored bars on the right-hand side of each sub-panel assign samples to wild (w) or domesticated (d) subpopulations according to the legend at the bottom right.

**Extended Data Figure 6:**
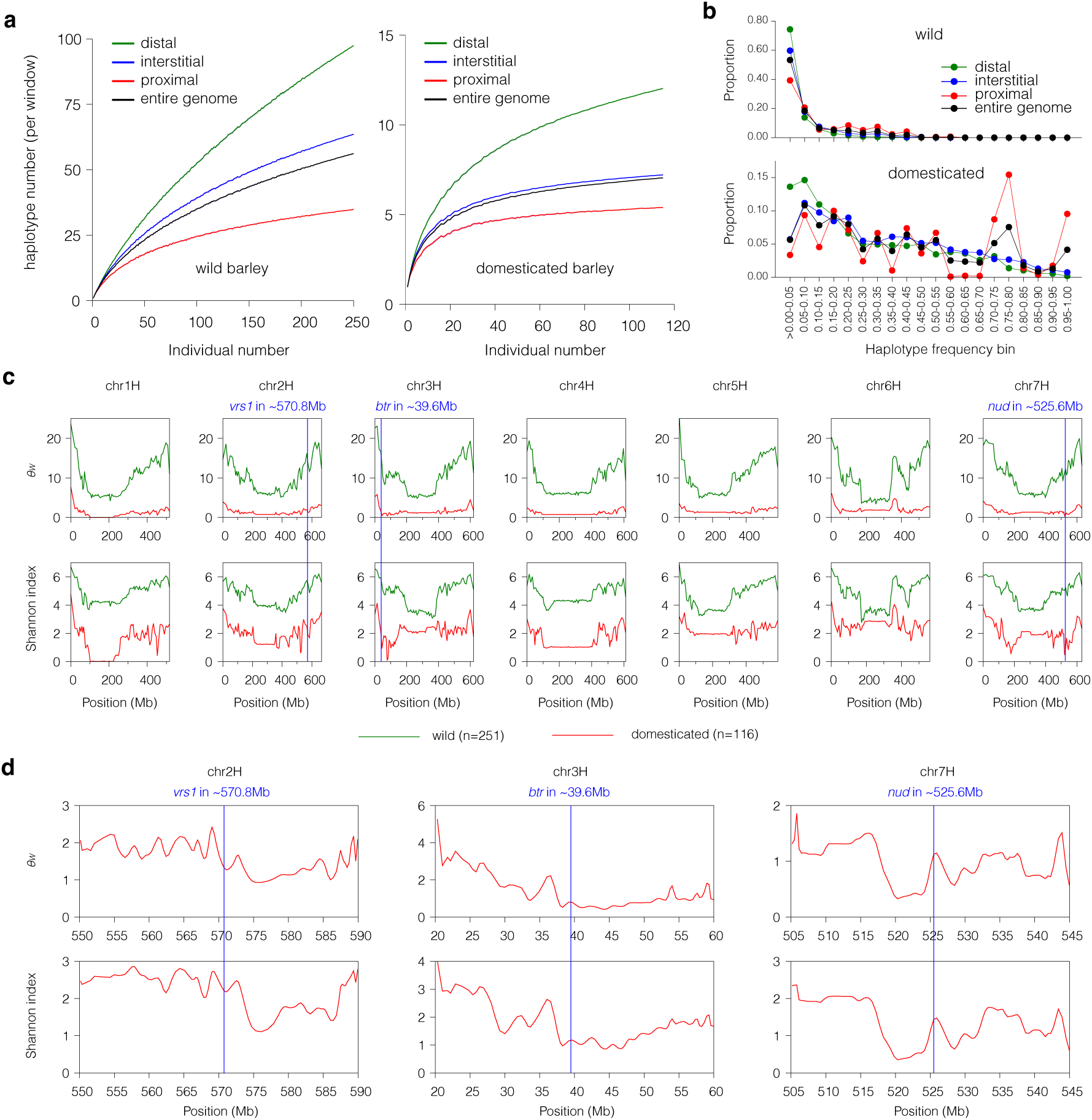
Haplotype-based diversity statistics. **(a)** Number of observed distinct haplotypes per genomic window in wild and domesticated barley as a function of sample size. The solid line and shaded area represent, respectively, the average and 95% confidence interval of 100 random sub-samples. **(b)** Haplotype frequency spectra (bin size: 0.05) in wild and domesticated barley in different genomic compartments. **(c)** Watterson’s θ and the Shannon index along the seven chromosomes of barley in wild and domesticated forms. Blue lines mark the location of *Btr1/2*, *Vrs1* and *Nud* loci. **(d)** Haplotype diversity (θ_W_ and Shannon index) in domesticated barley around these loci.

**Extended Data Figure 7:**
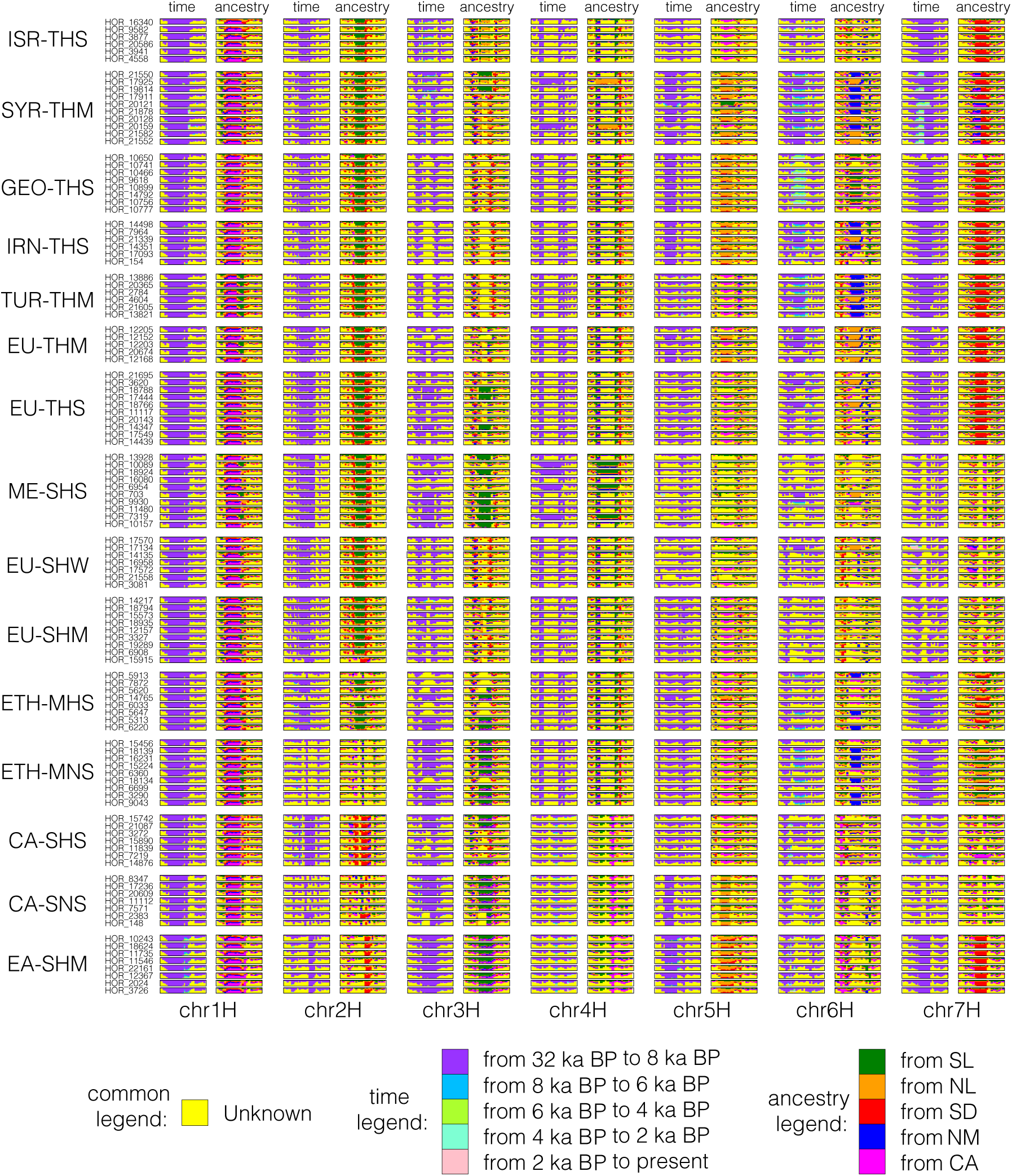
Spatiotemporal origins of haplotypes in domesticated barley. The inferred times at which haplotypes entered the domesticated gene pool and the most likely wild source populations are shown along the genome (20 Mb windows) for 116 domesticated barleys from 15 populations. Colors correspond to periods (time) and population (ancestry) as indicated in the legend. Yellow color indicates unknown origins.

**Extended Data Figure 8:**
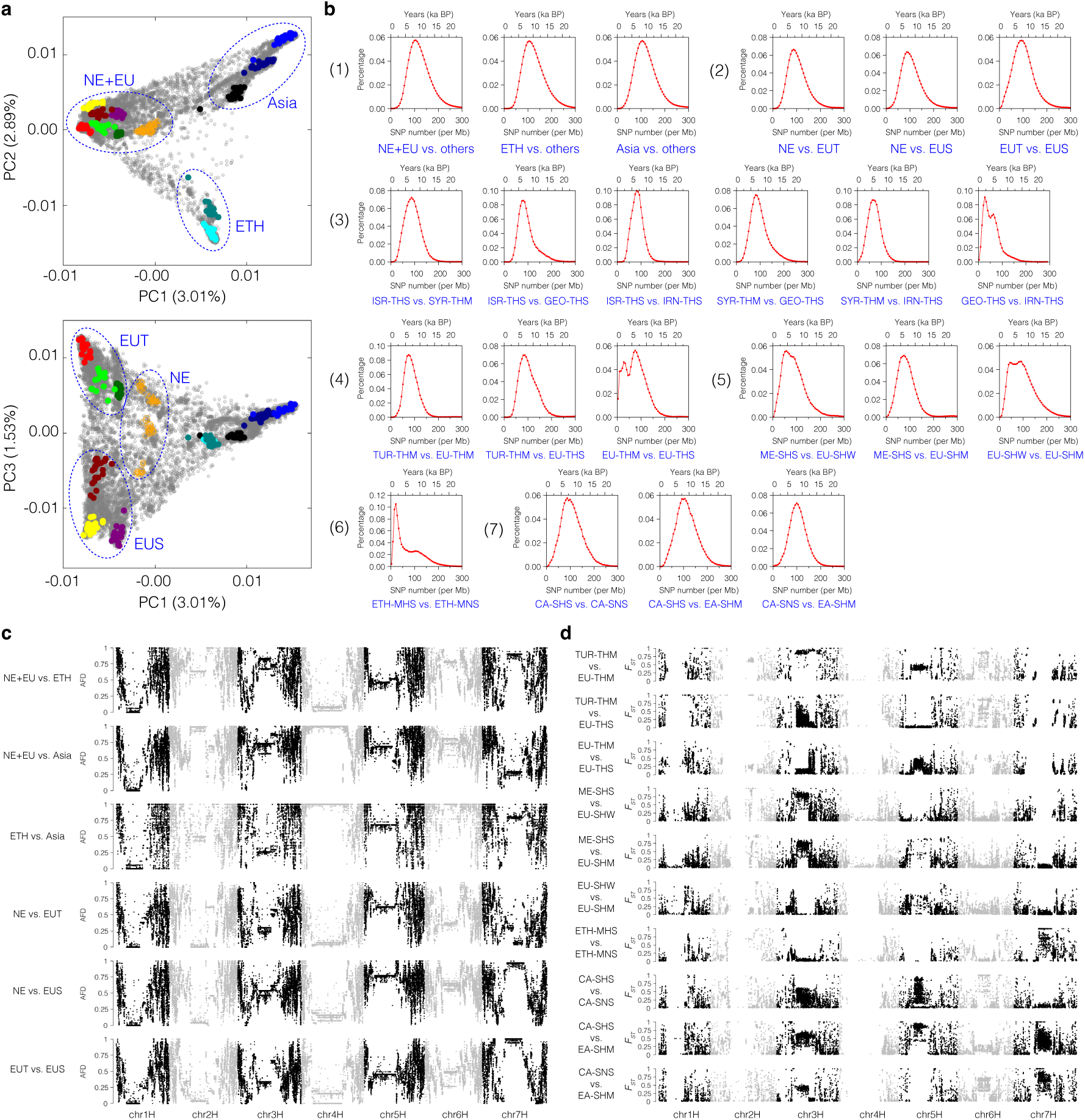
Divergence of domesticated barley populations. **(a)** Principal component analysis of 19,778 based on genotyping-by-sequencing data of Milner et al.^26^ (62,888 biallelic SNPs). Samples analyzed in this study are shown in non-gray color. Blue circles delineate the groups used for the comparisons in panel **(b)**: NE – Near East; EU – Europe and Mediterranean Basin; ETH – Ethiopia; Asia – Central and East Asia; EUT – EU two-rowed; EUS – EU six-rowed. **(b)** Distribution of sequence divergence between populations of domesticated barley. The comparisons are indicated in blue font below the sub-panels. **(c)** absolute allele frequency difference (AFD) between different domesticated barley populations in sliding windows (size: 100 kb, shift: 20 kb) along the genome. AFD was computed on the haplotype matrix of high-coverage (∼10x) samples **(d)** F_ST_ in sliding windows (size: 100 kb, shift: 20 kb) along the genome. F_ST_ was computed from the SNP matrix of all samples (matrix SNP2, see **Methods**).

**Extended Data Figure 9:**
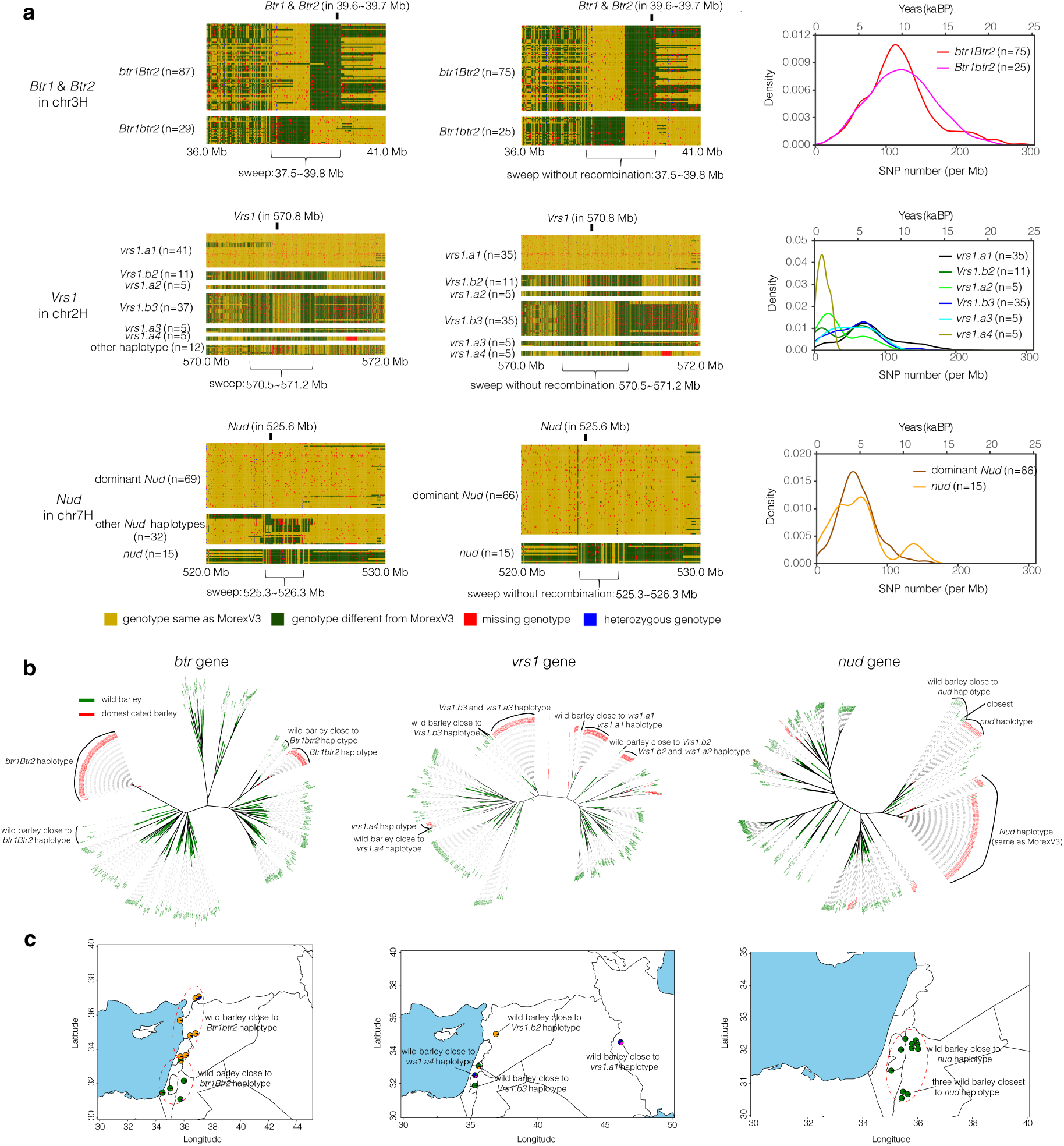
Origins of haplotypes of domestication genes. **(a)** The left-hand panels show the SNP haplotypes at the *Btr1/2*, *Vrs1* and *Nud* loci. The middle panels show the SNP haplotypes only of samples without recombination events that most likely occurred post-domestication. The right-hand panel shows the distribution of sequence divergence (SNPs per Mb) at these three loci. Only samples without recombination events were used. The window size for computing SNP numbers was 100 kb (shift: 20k kb). **(b)** Neighbor-joining tree of wild and domesticated barley computed from SNPs in the “sweep” interval denoted in panel **(a)**. **(c)** Collection sites of those wild barley samples whose haplotypes at the three loci are most closely related to domesticated haplotypes.

**Extended Data Figure 10:**
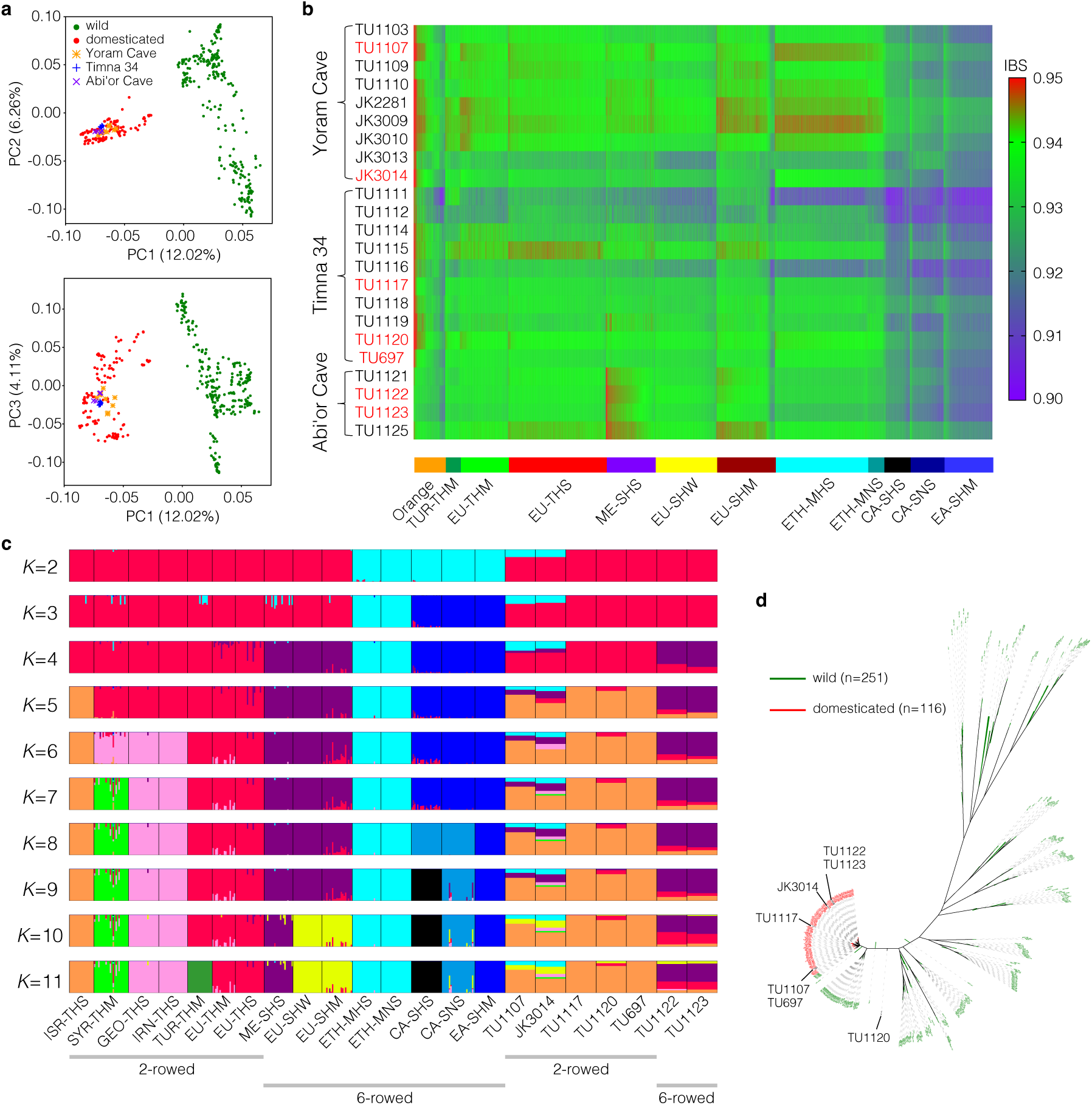
Ancient samples in the diversity space of extant barleys. **(a)** PCA on high-coverage wild and domesticated samples onto which 23 ancient barleys were projected. **(b)** Heatmap showing the identity-by-state (IBS) similarity between 23 ancient samples and 19,778 domesticated samples genotyped by Milner et al.^26^. Red font marks ancient samples sequenced at high coverage. Ancient samples with high similar to the “Orange” and ME-SHS population. The “Orange” population included Western Asian samples. **(c)** Ancestry coefficients as determined by ADMIXTURE with the number ancestral population (K) ranging from 2 to 11. The SNP matrix for the ADMIXTURE runs included 302 domesticated samples and high-coverage ancient samples. The widths of the bars corresponding to individual ancient samples have been increased by a factor of 20. (**d)** Neighbor-joining tree of 251 wild barleys, 116 domesticated barley and 7 high coverage ancient barleys. The tree was computed from 1.75 M biallelic SNPs in the interval 150 to 200 Mb on chromosome 1H. No MAF filter was applied.

